# 3D-super-enhancers are condensate-associated cis-regulatory communities

**DOI:** 10.1101/2024.02.01.578210

**Authors:** Jie Lv, Kelsey A. Maher, Li Dong, Virginia Valentine, Seth Staller, Alaguraj Veluchamy, Li Tian, Yuna Kim, Bensheng Ju, Marcus Valentine, John Easton, Stanley B. Pounds, Steven Burden, Brian J. Abraham

## Abstract

Transcription proteins are concentrated at nuclear transcriptional condensates. These condensates contain cis-regulatory elements (CREs), including enhancers and promoters, that are thought to regulate genes in the same condensate. The roles of condensates are of great current interest, but research into their function is limited by an inability to comprehensively identify their associated CREs. Here, we present a conceptual framework and algorithm, BOUQUET, for integrating genome topology, chromatin occupancy, and graph theory to associate CREs and transcription protein machinery with target genes and identify exceptionally protein-rich communities that interact with condensates. BOUQUET uncovers surprising quantitative correlations between community protein accumulation and gene expression phenotypes by combining accurate CRE-gene assignment with co-activator binding profiles. A small subset of communities, which we call “3D-super-enhancers,” is exceptionally protein-rich. BOUQUET-predicted 3D-SEs are comparable in number to co-activator nuclear puncta, and all genes known to interact with co-activator condensates in embryonic stem cells are within 3D-SEs. 3D-SEs are enriched for association with cell identity genes across mammalian tissues. Microscopy analyses show frequent co-localization and co-expression of genes from the same 3D-SE within a single co-activator punctum, suggesting 3D-SE components interact with co-activator condensates. Thus 3D-SEs correspond to co-activator puncta, which nominates additional condensate-associated genes and CREs.

## INTRODUCTION

Gene transcription is regulated through physical interactions among cis-regulatory elements (CREs) [e.g. enhancers and promoters], their bound proteins [e.g. transcription factors (TFs), co-activators, and RNA Polymerase II (Pol II)], and one or more target genes. Several critical transcription-regulating proteins have been observed to form nuclear bodies alternately called clusters, foci, droplets, puncta, or condensates, suggesting that dense nuclear protein accumulations are mechanistically critical for the regulation of transcription (1–3). For brevity, we use the term “condensate” and intend it to mean protein-dense, membraneless, nuclear puncta with internal environments that are distinct from the surrounding nucleoplasm, though we recognize the importance of ongoing work that probes the biophysical mechanisms of these partially divergent models. Condensates can contain high concentrations of a variety of biomolecules, including Pol II, TFs, cofactors, and chromatin regions that each facilitate gene control. However, precisely which genes associate with which types of condensate, how protein-bound chromatin regions interact with condensates, and how transcriptional products are produced by condensates remain obscured. To date, individual condensates and the genes that physically associate with them have been primarily examined using single-gene microscopy case studies (4–12). Here, we propose that an approach utilizing integrative omics analyses can identify distinct sets of spatially interacting, protein-bound CREs that elucidate the transcriptional regulatory relationships among chromatin regions and their bound protein apparatus.

The few genes that have been experimentally verified to associate with transcriptional condensates are genomically proximal to domains with very high levels of co-activator binding, suggesting that accumulation of co-activator signal at relevant CREs has some degree of correlation to nuclear co-activator puncta. (4–7, 9). To this end, co-activator ChIP-seq has been used to identify super-enhancers (SEs), i.e., linearly assembled enhancer sets with exceptionally high co-activator signal, but SEs are only somewhat informative for detecting condensate-associated chromatin regions (13). Among their drawbacks, SEs only assemble genomically proximal enhancers, and these assembled enhancer domains are assigned only to their single most proximal gene (13–15). This process fails to incorporate the important regulatory contributions of genome organization on gene expression, including long-distance enhancer-promoter interactions, insulated chromatin loops (e.g. insulated neighborhoods) that constrain and/or facilitate such interactions, the ability of other types of CREs to act as enhancers, or the complexity of densely interconnected gene-CRE regulatory networks (e.g. chromatin hubs, clusters, or cliques) (16–38). Further, when studied in mESCs, the number of linear SEs differs from the number of observed co-activator puncta by several fold (4, 13, 14, 39). Thus, even if assuming all SEs form co-activator condensates, the genomic elements affiliated with the majority of co-activator condensates remain unknown.

In this study, we used co-activator binding profiles and genome topology measurements to predict the genes and CREs associated with transcriptional protein condensates. To accomplish this goal, we built BOUQUET, a computational pipeline that leverages a label-propagation machine learning method from graph theory to uncover the regulatory networks of transcriptional co-activator–bound CREs, or “communities”. We find that, in single cells, genes within the same community have elevated degrees of co-expression, and CREs within the same community have correlated accessibility, consistent with these elements occupying a shared sub-nuclear environment. We observe that, in mESCs, the distributions of loading for Mediator subunits, BRD4, and other transcriptional proteins are asymmetric across communities and their summed levels significantly correlate with gene expression. Community composition reliably predicts the transcriptional impact following a variety of perturbations, including the deletion of constituent CREs and the depletion of a co-activator. We identify a subset of communities with exceptional co-activator signal and call them “3D super-enhancers”, or “3D-SEs.” 3D-SEs are enriched for genes that control specific cell identities across a variety of cell types. Extremely high-signal CRE assemblies, such as those at the histone-encoding gene loci that form the histone locus body condensate, are also 3D-SEs, despite being undervalued by current approaches. Finally, we confirm via fluorescence microscopy that *Sptlc2*, which shares a 3D-SE with the cell identity gene *Esrrb*, also shares its co-activator condensate; these genes are co-regulated, despite their sizable genomic separation. Together, our approach sheds light on the genes that associate with spatial concentrations of co-activators and the genetic elements that regulate them, opening new lines of investigation into the functions of transcriptional condensates.

## MATERIALS AND METHODS

### Identification of cis-regulatory elements (CREs)

In this study, we examine cis-regulatory elements (CREs), specifically genetic promoter and enhancer elements. We defined the promoter set for each gene as the 4 kb regions centered on each of its transcripts’ transcription start sites (TSS). For our reference gene annotation, we downloaded RefSeq Release 108 for GRCm38.p6 from the NCBI FTP site, https://ftp.ncbi.nlm.nih.gov/refseq/M_musculus/annotation_releases/108/GCF_000001635.26_GRCm38.p6/GCF_000001635.26_GRCm38.p6_genomic.gtf.gz. In order to focus on high-confidence gene annotations, we filtered 1) *against* genes annotated as “unknown transcript,” 2) *against* MGI-predicted genes, i.e. those whose gene names start “Gm,” and 3) *for* genes whose identifiers start with “NM” or “NR”, which are curated protein-coding and non-protein-coding genes, respectively, resulting in 40,618 genes for downstream analysis. We measured the activity of each promoter by the number of H3K27ac ChIP-seq (GSM1526287) reads mapped to the region with at least one base pair overlap using bedtools intersect. Then we defined active genes as those with a TPM>=1 and with at least one promoter ranking in the top two-thirds of promoters as measured by H3K27ac coverage. For expressed genes/promoters with multiple transcript isoforms, we collapsed all promoters for a given gene using bedtools merge. We defined genetic enhancers by calling peaks using H3K27ac ChIP-seq data via the SEASEQ pipeline (v3.0) (40). Briefly, reads were mapped to the mouse reference genome sequence version mm10 (GRCm38). To identify narrow enriched regions (peaks) for H3K27ac, MACS (v1.4.2) (41) was utilized through SEASEQ with the following parameters: keep-dup= auto, band width = 300, model fold = 10,30, p-value cutoff = 1.00e-09. If corresponding control libraries existed, they were used in peak-calling. Any H3K27ac ChIP-seq peak that did not overlap with a promoter region was considered an enhancer. Collectively, the promoter and enhancer elements as defined above are referred to in this study as CREs.

### Building Optimized Units of QUantified Enhancer Topologies (BOUQUET): inferring structure-informed gene regulatory communities

We built enhancer/promoter networks in mESCs by combining a reference gene list, an active enhancer map, an active promoter map, H3K27ac HiChIP loops, Insulated Neighborhoods (INs), and, optionally, topologically associating domains (TADs) (42).

#### Step 1: Building network of CREs

Nodes of the network, i.e. CREs, were defined using H3K27ac ChIP-seq and a reference gene annotation. Specifically, CRE nodes were the collapsed union of ChIP-seq-defined peaks and active gene promoters as defined above. Briefly, 11,051 collapsed regions that overlapped a promoter by >=1 bp were identified as promoter nodes, and H3K27ac ChIP-seq peaks that did not overlap at all were considered enhancer nodes. We added an edge between two nodes if they fit either or both of two criteria: 1) both nodes overlapped with the two anchors of the same HiChIP loop, defined above; 2) an enhancer node and a promoter node were both fully contained within the smallest IN, i.e. minimal number of bps, that contained the promoter (Supp. Data S1-S4). Our study prioritized the characterization of regulatory communities governing active gene expression, but regions containing promoters of lowly or non-expressed genes are also accounted for in our workflow both explicitly and implicitly. Of the 2,394 loop-involved bins that overlapped promoter regions of lowly or non-expressed genes (TPM ≤ 1), 1,914 (80%) also either overlapped an H3K27ac ChIP-seq peak or the promoter of an expressed gene, and are thereby included in at least one community. While 1,561 individual silent gene promoters meet these inclusion criteria, they represent a minority (12%) of all silent promoters genome-wide (12,522 total), consistent with the biological expectation that high-confidence regulatory loops preferentially but not exclusively connect transcriptionally active loci.

#### Step 2: Identifying communities of CREs

With the CRE network as input, we employed the R package igraph (v.1.2.6) (43) to infer communities. Communities are defined in graph theory as sets of nodes that are highly interconnected, i.e., many edges connect nodes within a community, and few edges connect nodes from distinct communities (44). We used a modified Label Propagation (LP) algorithm with back-filling to detect communities of CREs for each expressed gene (45). We first identified subsets of nodes in the genome-wide network that are completely disconnected from others, i.e., network components. For a given gene within a component, the gene node and its neighbors, i.e., the CREs connected to it directly by loops or by INs, was initialized with the same community label whereas every other node in the component was initialized with a unique community label. At every iteration of propagation, each node updates its community label to the one that occurs with the highest frequency among its directly connected neighbors. The LP algorithm resolves when every node within a component has the label that the maximum number of its neighbors also have. As labels propagate through the component, a single label can quickly become dominant in a densely connected group of nodes but will have trouble crossing a sparsely connected region of the component. As a result, densely connected groups of nodes quickly reach a consensus on a unique label. At resolution those nodes that are assigned the same label of the given gene are considered part of that gene’s community. We appended a step to this established LP approach to ensure that directly linked nodes were captured; we used a back-filling method to rescue any directly linked nodes, i.e. CREs that loop to the promoter or that are within the same smallest IN as the promoter, that did not share the same community label of the given gene after LP process. Consistent with the graph theory literature, we refer to the highly interconnected subsets of nodes produced by this method as “communities” (Supp. Data S5).

#### Step 3: Calling 3D-Ses

We extracted 3D-SEs from all communities according to their constituent CREs’ collective transcriptional protein coverage, which was calculated separately for each mark we studied, including co-activators, H3K27ac, CTCF, RONIN, etc. For each community and for each mark, we summed the relevant ChIP-seq reads across all its constituent CREs as its measurement of regulatory signal loading. When ranking communities by their regulatory signal loading, we observed a distribution where a limited number of communities have exceptionally high cumulative signal. We identified the point in the distribution at which the line y=x is tangent to the curve and used this point as a cutoff as described previously (13). Communities above this signal cutoff were defined as 3D-SE CRE communities (3D-SEs) (Supp. Data S6).

#### Step 4: Optional step for expanded CRE community assignment

Some researchers may be interested in studying the regulatory communities of CREs that are not covered by direct loops or by INs. If the optional “expanded CRE community assignment” feature is used, following the completion of the community detection process described above, any remaining unassigned, active enhancers that are within a TAD are assigned using a combined linear proximity/TAD approach. That is, the linearly most proximal promoters in both the upstream and downstream directions are considered for each enhancer, and the enhancer is assigned to the most linearly proximal promoter within the same TAD, if any such promoter exists, using bedtools closest. TADs can constrain CRE interactions, potentially improving the accuracy of the linear proximity-focused, one-to-one assignment of enhancers to genes. We include this step as an opt-in assignment feature to facilitate near complete level of CRE community assignment, but it has lower precision than loop/IN-informed CRE connections (Supp. Fig. 1A) (42, 46–49).

### BOUQUET-lite

Though its most obvious use case is assessing genome looping, H3K27ac HiChIP data implicitly measures proxies of additional gene regulatory properties, including CRE activity and gene expression. Thus, we believed H3K27ac HiChIP alone could be used to enable CRE community detection. We designed a version of BOUQUET, BOUQUET-lite, that uses only H3K27ac HiChIP, which enabled us to study CRE communities in additional mouse tissues where only H3K27ac HiChIP data were readily available. In case an enhancer map is not available in the matched cell type, an alternative method is available to infer candidate loop anchors using H3K27ac HiChIP itself. We used HiChIP-Peaks (v0.1.2) with “-f 0.4” to call enriched peak regions from H3K27ac HiChIP datasets and used the resulting peak set as active CREs. In parallel, promoters (TSS +/-2kb) were ranked by their coverage of H3K27ac HiChIP reads calculated using bedtools intersect, and the top two-thirds were retained to represent expressed genes. H3K27ac HiChIP peaks and expressed promoters identified thusly were collapsed using bedtools merge to generate a set of candidate loop anchors, which were used as input for Hi-LOW as above. In case INs are not available, we relied strictly on H3K27ac HiChIP loops to infer CRE connections and build the initial network of CREs genome-wide (Supp. Data S7-S9). Finally, H3K27ac HiChIP read coverage was used to rank each community instead of co-factor ChIP signal.

These resulting loops, H3K27ac HiChIP peaks, and H3K27ac HiChIP coverage BAMs were used as input for BOUQUET-lite in the following cell/tissue types: mouse embryonic stem cells (GSE99519), thymocytes (GSM5268039), SST-neurons (GSM4551965), regulatory T cells (Treg) (GSM3059349), and cardiomyocytes (GSM6916796) (Supp. Data S10-S14). A cutoff for 3D-SEs was similarly defined as described above (Supp. Data S15).

### CRE accessibility correlation – LASER

To calculate CRE accessibility correlation, we created a customized CRE-by-cell matrix. We quantified the accessibility of each CRE in each cell using the function FeatureMatrix() of the Signac package, using the fragment file and the CRE node list defined above as input. CRE read counts were normalized by the ‘LogNormalize’ method using the function NormalizeData().

To measure the correlation of pairwise CRE accessibility, we binarized (set CRE read counts to 0 or 1) the CRE-by-cell matrix and calculated the Odds Ratio (OR) using the 2 x 2 read count table for each pair of CREs. To test the statistical significance of CRE ORs between different groups, we used a bootstrapping technique to generate simulated CRE-by-cell matrices. This was done by randomly resampling cells with replacement, and it allowed us to compute the following statistic from each simulated matrix: the difference of median ORs between two groups, e.g., community vs. background. Since this statistic largely follows a normal distribution (with only 8 out of 190 bootstrap distributions showing significant p-values (<0.05) by the Shapiro-Wilk test across bootstrap simulations), we calculated a p-value while assuming the null hypotheses, i.e. that there is no difference of OR between the two groups compared. We developed a method, Loci ATAC-Seq Enrichment Resampling (LASER), to perform the process described above. The LASER method is estimated to run in time O(n^2^) (with n being the number of total CREs), making the method computationally intractable for calculating all genome wide CRE pairs at once. To overcome this limitation, LASER splits the genome into pairs of chromosomes (190 pairs of 20 mouse chromosomes) and calculated correlations between peaks on all pairs of chromosomes in parallel. This method substantially reduced runtime and made it feasible to calculate pairwise CRE accessibility correlation genome wide.

See Supplementary Material for detailed materials and methods.

## RESULTS

### Assembling CRE communities using BOUQUET

We developed “BOUQUET” (Building Optimized Units of QUantified Enhancer Topologies), a computational pipeline that uncovers communities of co-activator–bound CREs based on genome-wide CRE interactions (Fig. 1A, Supp. Fig. 1A, Methods) (17, 31). Our fundamental hypothesis is that accurate assignment of transcription apparatus using genome organization will aid the identification of collections of especially-protein-rich genome regions that are associated with nuclear condensates. To accomplish this goal, BOUQUET connects CREs into networks using high-confidence topology measurements, uses a label-propagation machine learning algorithm from graph theory to subdivide CRE networks into highly interconnected sets called “communities,” and quantifies these communities based on their summed transcription protein signal at their constituent CREs (Methods, Supp. Data S1-S5). BOUQUET begins by considering a pair of CREs to be connected if they meet at least one of two criteria: they are connected by a high-confidence HiChIP loop; or they are contained within the same Insulated Neighborhood (IN) (see Methods). Insulated Neighborhoods (INs) are chromatin loops formed by contacts between CTCF/cohesin-bound regions whose disruption can impact the regulation of transcriptional bursting by chromatin condensates (9, 35–37). Our use of IN-derived connections is supported by the similar read counts observed for CRE-pairs connected by high-confidence loops versus CRE-pairs connected by INs but not high-confidence loops when such CRE-pairs are separated by similar distances (Supp. Fig. 1B) (50). Using only high-confidence HiChIP loops in mouse embryonic stem cells (mESCs), only 74.8% of the active enhancers and 81.3% of the active promoters in mESCs were directly connected to another activating CRE; further, not every high-confidence loop connected pairs of activating CREs (Supp. Fig. 1C, D). When we combined high-confidence H3K27ac HiChIP loops and INs, we could link 89.9% of active enhancers and 91.9% of active promoters to at least one other CRE, giving us a near-comprehensive coverage of activating CREs by genome topology (Supp. Fig. 1C). We tested multiple thresholds for high-confidence loops, and, despite the expected variation in loop numbers, promoter and enhancer coverage remained relatively stable (Supp. Fig. 1E).

**Figure 1:**
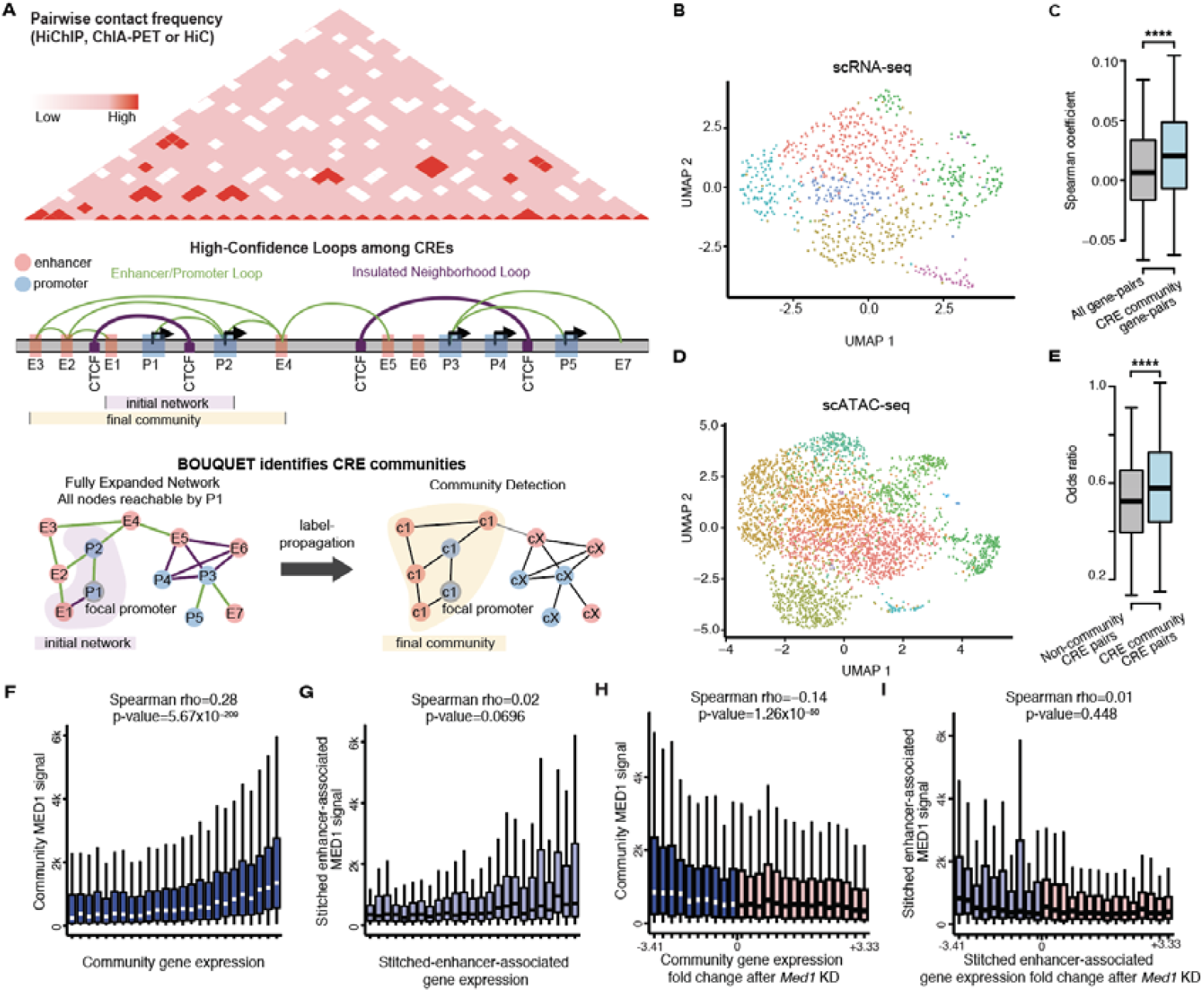
BOUQUET identifies cis-regulatory element communities and uncovers relationships between accumulated protein apparatus and gene expression. **(A)** Workflow of Building Optimized Units of Quantified Enhancer Topologies (BOUQUET). High-confidence loops (green arcs) connecting pairs of cis-regulatory elements (CREs; promoters in blue, enhancers in red) are identified by H3K27ac HiChIP using the Hi-LOW pipeline (see Supplemental Methods). Interactions between pairs of CTCF binding sites that are also bound by cohesin form Insulated Neighborhoods (IN) (purple arcs). Both high-confidence H3K27ac HiChIP loops and INs are used by BOUQUET to connect CRE-pairs. CRE-pairs are assembled into an initial CRE network (violet cloud). Using label-propagation, BOUQUET refines the initial network to identify a focal gene’s final CRE community by uncovering densely interconnected subnetworks (yellow cloud). See Methods for additional details. **(B)** UMAP plot of scRNA-seq from 847 mESCs. Each point represents an individual cell, with colors indicating Seurat-derived clusters. **(C)** Distributions of Spearman’s correlation coefficients from scRNA-seq data across 847 mESCs, comparing co-expression of (i) genome-wide control gene pairs (gray; median r= 0.0065) and (ii) gene-pairs within the same CRE community (blue; median r=0.02). p-value < 2.0×10^-308^ (Two-sided Wilcoxon Rank Sum Test; see Supplemental Methods). Whiskers represent interquartile range. (**D**) UMAP plot of scATAC-seq from 3,171 mESCs. Each point represents an individual cell, with colors indicating Seurat-derived clusters. (**E**) Distributions of odds ratios (ORs) from scATAC-seq data across 3,171 mESCs, comparing co-accessibility of (i) genome-wide control CRE pairs (not within the same CRE community; gray; median OR=0.52) and (ii) CRE-pairs within the same CRE community (blue; median OR=0.58). p-value =1.6×10^-25^ (bootstrapping procedure assuming normal distribution). Whisker length represents interquartile range. (**F**) Distributions of community-assigned per-gene MED1 signal across grouped genes. Genes are sorted into 25 groups by increasing expression level. (**G**) Distributions of ROSE-assigned per-stitched-enhancer-associated gene MED1 signal, arranged into 25 groups by increasing gene expression level. (**H**) Distributions of community-assigned per-gene MED1 signal across grouped genes following knockdown of *Med1*. Genes are sorted into 25 groups by their expression fold-change after *Med1* knock-down (KD). Blue indicates downregulation of community gene expression following *Med1* KD; pink indicates upregulation of community gene expression following *Med1* KD. (**I**) Distributions of ROSE-assigned per-stitched-enhancer-associated gene MED1 signal following knock-down of *Med1*. Genes are sorted into 25 groups by their expression fold-change after *Med1* knock-down. Blue indicates downregulation of stitched enhancer-associated gene expression following *Med1* KD; pink indicates upregulation of stitched enhancer-associated gene expression following *Med1* KD.

We applied BOUQUET to identify CRE communities in mESCs, a well-established system for the study of co-activator condensates and their associated genes (4–7). For 23,084 active genes in mESCs, we identified 11,051 communities, with a median of 7 CREs per community. There was an overall positive correlation (Spearman rho = 0.46, p-value = 1.6 x 10^-245^) between the numbers of promoters and enhancers within a community, but a few showed a bias toward either promoters or enhancers (Supp. Fig. 1F). We next quantified ChIP-seq of MED1 at communities as an initial representative co-activator because of its established association with large CRE collections and transcriptional condensates, its mechanistic role in the facilitation of physical connectivity of TF-bound enhancers to promoters, and its regulation of Pol II elongation (51). Several properties of CRE communities are highly correlated, e.g., communities with greater numbers of CREs tended to also have higher MED1 signal, greater numbers of overall interactions (i.e. “edges”), and greater number of interactions from the community’s focal gene (Supp. Fig. 1G, Methods) (16).

### Elements of the same community show elevated activity correlation

Having identified CRE communities genome-wide, we next investigated the properties of CREs assigned to the same community. If the CREs within communities are functionally linked by sharing the same sub-nuclear environment, we would expect that elements of the same community would demonstrate higher levels of coordination in their activities, i.e. gene expression and CRE activity. To test this hypothesis, we generated scRNA-seq in mESCs. Though clusters were algorithmically identified, the whole sample showed overall homogeneity in expression profiles, which supported our use of each cell as a replicate (Fig. 1B). For each pair of expressed genes (gene-pair), we calculated Spearman’s correlation coefficients for their expression levels across all cells (Fig. 1C). For gene-pairs that were not part of the same community, the median expression correlation level was low (r=0.0065). In contrast, gene-pairs within the same community showed a two-fold higher median expression correlation level (r=0.02) and a significant but modestly higher trend of co-expression than non-community gene-pairs (p = 2.23×10^-308^, two-sided Wilcoxon Rank Sum Test). The statistically significant correlation between shared community membership and gene co-expression held across eight other commonly used statistical tests of correlation (Supp. Fig. 1H). This finding supports the hypothesis that gene expression is more tightly coordinated among genes that share a community than between genes that do not, but this correlation is modest, likely due to the complex set of processes that contribute to expression.

To validate the relevance of connections derived from INs in BOUQUET, we built an additional set of CRE communities using loops alone and compared co-expression of gene-pairs from these communities to our index community set (loops + INs). Community gene pairs identified using both approaches exhibited comparable levels of expression correlation, with Spearman correlation coefficients of r=0.021 for the loops-only approach and r = 0.020 for the index approach (Supp. Fig. 1I). Most community gene pairs were recovered by both approaches, while only a minority are uniquely detected by either method (Supp. Fig. 1J). Community gene-pairs shared between both approaches exhibited the highest expression correlation, whereas gene-pairs unique to either loop+IN or loops-only communities shows statistically indistinguishable correlation levels (p=0.16, two-sided Wilcoxon rank-sum test; Supp. Fig. 1K). These results support incorporating INs as additional evidence for defining CRE communities.

To further examine the degree of coordinated CRE activity within communities, we performed single-cell ATAC-seq (scATAC-seq) in mESCs. ATAC-seq measures chromatin accessibility to the Tn5 transposase, which is an indicator of a region’s ability to bind to transcription-regulating proteins that may be used as a proxy for enhancer activity (52–54). UMAP clustering analysis of our scATAC-seq data reiterated that our cell population is generally homogeneous, here at the chromatin accessibility level, supporting our handling of cells as replicates (Fig. 1D). In contrast to scRNA-seq, there is an upper limit on the number of ATAC-seq integration sites across two copies of the same CRE, which compresses the possible dynamic range of measurements at a given locus (52). To perform robust statistical analyses of co-accessibility, we developed an algorithm that overcomes this range limitation, Loci ATAC-Seq Enrichment Resampling or “LASER,” which calculates odds-ratio-based measurements of the co-accessibility of CRE-pairs across a set of individual cells. We used LASER to establish a background CRE co-accessibility distribution against which to evaluate the co-accessibility of CREs within a single community by randomly selecting pairs of CREs not from the same community. We compared the co-accessibility of these randomized CRE-pairs with pairs of CREs within the same community (Fig. 1E). CRE-pairs within the same community showed a significant but modestly higher trend of co-accessibility than randomized non-community CRE-pairs (median Odds Ratio = 0.58 vs. 0.52, respectively; p=1.6×10^-25^, bootstrapping procedure). Altogether, these results suggest that activities of within-community CREs are correlated, but these correlations are modest, likely due to the influences of additional gene regulatory processes.

### BOUQUET communities recapture CRE sets that regulate condensate-associated genes

We next probed the communities of known co-activator condensate-associated genes for confirmed and novel CRE relationships. One such gene of interest is *Sox2*, which encodes a master transcription factor that regulates pluripotency and associates with condensates of BRD4 and of MED1 in mESCs. (9, 55–60). Previous studies revealed that *Sox2* is regulated by both a proximal linear SE and a distal SE approximately 100 kb downstream, called the “Sox2 Control Region” (SCR) (Supp. Fig. 2A) (13, 59, 61). Further, three-way microscopy imaging previously revealed the SCR’s role in regulating transcriptional bursting of *Sox2* during interactions among *Sox2*, the SCR, and transcriptional condensates (9). Consistent with this work, our pipeline highlighted interactions between *Sox2*’s promoter and both the proximal linear SE and the SCR, and that all of these elements were within the same CRE community (Supp. Fig. 2A, B). Further, BOUQUET’s prediction that *Sox2* is the only protein-coding gene in its community was validated by the previous observation that genetic deletion of the SCR in mESCs only significantly impacts the expression of *Sox2*, leaving the expression of surrounding, linearly proximal genes unaffected (59). Consistent with the association of *Sox2*, the SCR, and co-activator condensates, *Sox2*’s community was in the top 1% most MED1-rich and top 2% most BRD4-rich (Supp. Data S5).

A second gene of interest is *D7Ertd143e*, a mammal-specific polycistronic pri-microRNA, whose mature miRNAs, *miR290* through *miR295*, regulate ESC pluripotency. *D7Ertd143e* has been shown to associate with nuclear co-activator condensates of MED1 and of BRD4 and, correspondingly, the largest linear SE (Supp. Fig. 2C) (4, 62). Previous single-gene expression analysis showed that deleting a CRE within the *D7Ertd143e*-associated SE caused significant downregulation of *D7Ertd143e*. BOUQUET confirmed that the deleted CRE and *D7Ertd143e* were indeed within the same community in wild-type cells. We performed RNA-seq on CRE-deleted cells to study the effects of CRE deletion on other genes within the *D7Ertd143e* community (Supp. Fig. 2C, D, E, Supp. Data S16, S17) (63). As expected, we observed significant downregulation of *D7Ertd143e* in CRE-deleted cells (p-value=2.77×10^-107^) (63). Additionally, we found that the genes in the *D7Ertd143e* community were, on the whole, significantly more downregulated than control genes (p=5.9×10^-4^, Wilcoxon Rank Sum Test, two-sided) and that all but one expressed gene in the *D7Ertd143e* community were significantly downregulated (Supp. Fig. 2E, Supp. Data S17). Notably, expression was not diminished for genes immediately beyond the linear boundaries of the community, despite all of the community and linearly proximal genes being within the same TAD (Supp. Fig. 2E, Supp. Data S17). Consistent with the association of *D7Ertd143e*, its SE, and co-activator condensates, *Mir290’*s community is in the top 1% most MED1-rich or top 1% BRD4-rich (Supp. Data S5). These results suggest that large communities uncovered by our approach accurately predict the genes that will be affected by CRE perturbation.

### Community co-activator signal correlates with gene expression and expression response to co-activator perturbation

Previous CRISPR tiling screens showed that the effect of perturbing a given CRE on its target gene correlates with that CRE’s ChIP-seq signal (called “activity”) and its frequency of contacting the target gene (64). We therefore hypothesized that target gene expression should directly correlate with CRE signal, but this relationship would only be revealed if all relevant CREs were accurately associated with their target genes and their signal were combined. To test this hypothesis, we compared gene expression against BOUQUET-assigned co-activator signal or against genomically proximal co-activator signal using ROSE. The combined MED1 signal at each CRE community was significantly correlated with the expression of genes within that community by bulk RNA-seq in mESCs (Spearman rho=0.28, p-value=5.67×10^-209^) (Fig. 1F, Supp. Data S5); this relationship remained consistent across the different loop-calling thresholds we examined (Supp. Fig. 2F). In contrast, genomically linear enhancer/co-activator assignment showed much lower and insignificant correlation to expression levels of their assigned target genes (Spearman rho=0.02, p-value 0.0665) (Fig. 1G, Supp. Fig. 2G, Supp. Data S18). These results suggest both that accurately assigned co-activator signal better correlates with expression level and that topology-informed BOUQUET communities more accurately capture the co-activator–bound CREs relevant to target gene regulation.

If co-activator signal within a community is predictive of gene expression, then loss of that co-activator might more dramatically affect high-signal communities. To test this hypothesis, we acquired RNA-seq data from mESCs after *Med1* had been knocked-down (65). As hypothesized, there was a significant negative correlation between the expression response to *Med1* knock-down and MED1 signal assigned to a gene by BOUQUET; genes downregulated most profoundly were among the most MED1-rich communities, and vice versa (Spearman = -0.14, p-value = 1.26×10^-50^) (Fig. 1H, Supp. Data S19); this relationship was consistently observed across the different loop-calling thresholds we analysed. (Supp. Fig. 2H). In contrast, expression response to *Med1* knock-down showed no correlation with co-activator signal assigned by genomic proximity (Spearman = 0.01, p-value = 0.448) (Fig. 1I, Supp. Data S19). Thus, protein signal associated with genes using genome topology quantitatively predicts expression and response to perturbation.

### High correlation among activator marks at communities

Multiple transcriptional proteins form condensates and may be heterogeneously responsible for gene regulation by distinct CREs (4, 6–9, 66, 67). We therefore hypothesized that communities could be differentially enriched in different co-activators. We compared the genome-wide distributions of MED1, BRD4, CBP, or EP300 because each of these factors bind active CREs genome-wide, but they have distinct mechanisms of action. We confirmed that they have partially distinct binding sites across the genome but a high degree of correlation overall (Supp. Fig. 3A, B) (29). We also examined H3K27ac, a histone modification that marks nucleosomes adjacent to active CREs and that is commonly used for super-enhancer analysis (39, 68). The genome-wide distribution of H3K27ac was significantly correlated to that of the co-activators, making it an attractive alternative for studying systems where limited datasets are available (Supp. Fig. 3A, B). In contrast, other condensate-forming proteins (RONIN, CTCF) did not show similar degrees of correlation genome-wide (Supp. Fig. 3B) (6, 29, 69–71). The high correlation between activation marks was more pronounced when marks were compared using within-community signal (Supp. Fig. 3C, Supp. Data S5). Again, CTCF and RONIN did not show similar degrees of correlation at BOUQUET-derived communities, which we interpret to be due to their distinct preference for genomic regions (insulators and certain activating CREs for CTCF, housekeeping gene promoters for RONIN) (Supp. Fig. 3C, Supp. Data S5) (6, 29, 69, 70). Thus, despite some variation in each factor’s mechanism of action and chromatin binding at the individual-CRE level, co-activators are generally correlated in their collective accumulation at CRE communities.

Similar to MED1, we found that communities with higher accumulated levels of activation marks contained genes with higher levels of community gene expression (Fig. 2A). As seen for MED1, this correlation was stronger than the correlation of gene expression with linear-genome-based assignment of signal for the same factors (Supp. Fig. 3D, Supp. Data S20). Thus, community signal for co-activators and H3K27ac correlate highly with community gene expression.

**Figure 2:**
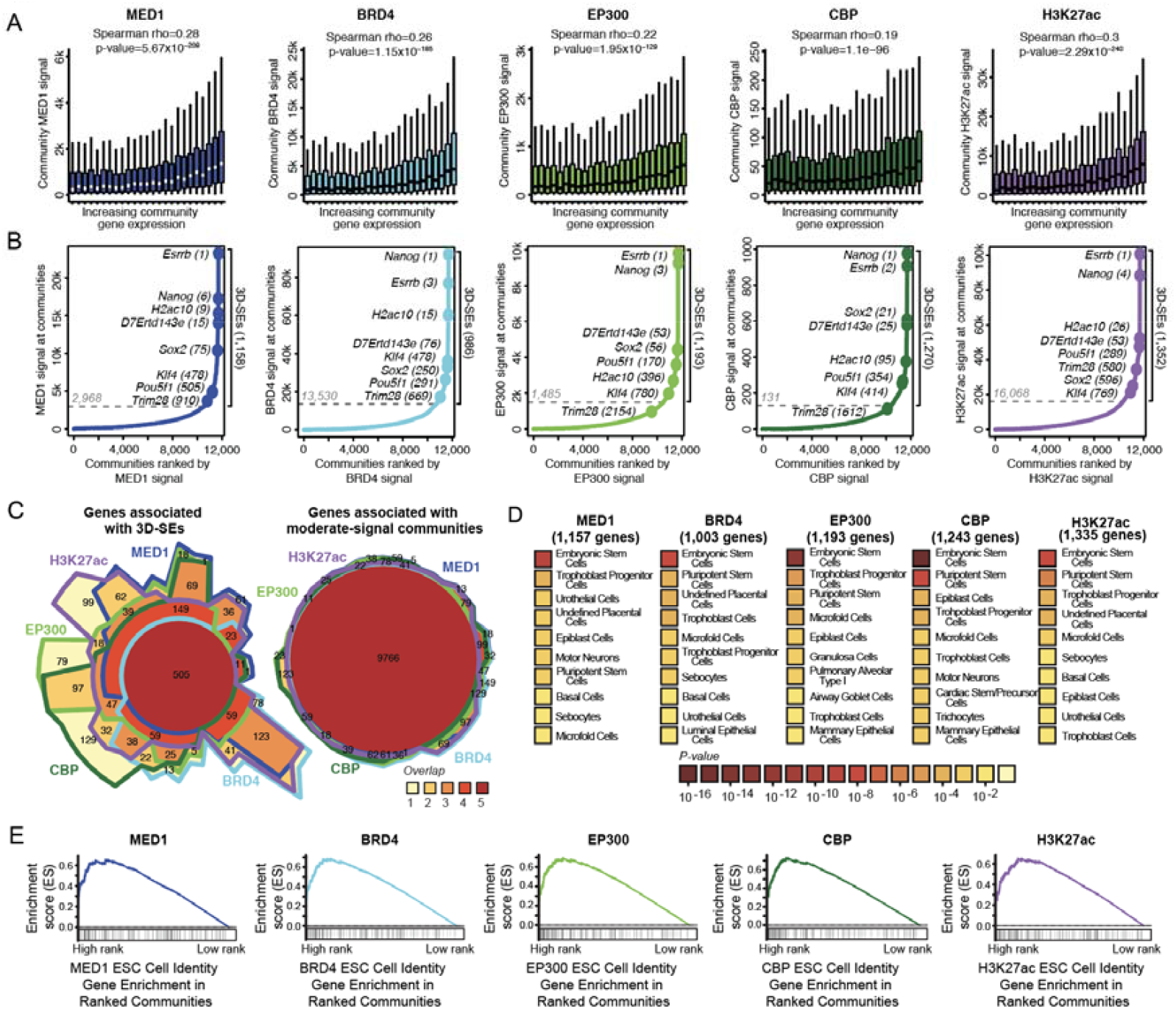
Ranking communities by co-activator binding reveals 3D super-enhancers. **(A)** Cumulative per-community ChIP-seq signal versus focal gene expression for MED1, BRD4, EP300, CBP, and H3K27ac. Communities are binned into 25 groups by increasing focal gene expression. Signal is quantified as the total mapped ChIP-seq reads across community CREs. The MED1 panel is duplicated from Fig. 1F and is provided here for ease of comparison. (**B**) Distribution of signal ranking across CRE communities for MED1, BRD4, EP300, CBP, and H3K27ac. X-axis: per-community signal quantified as summed ChIP-seq reads across community CREs; X-axis: relative community rank by per-community signal. Known condensate-associated genes are labeled. Total 3D-SE count is shown on the right; dashed line marks the algorithmic cutoff distinguishing 3D-SEs from moderate-signal communities (see Methods for additional details). (**C**) Chow-Ruskey diagrams showing the overlap of genes associated with 3D-SEs (left) and moderate-signal communities (right) for four co-activators and H3K27ac. The area of each section of the diagram is proportional to the number of genes within it. (**D**) Gene ontology analysis for ‘cell type’ term enrichment in 3D-SE genes ranked by one of four co-activators or H3K27ac. The top 10 most enriched terms from the PanglaoDB_Augmented_2021 database are shown for each factor (see Supplemental Methods for additional details). (**E**) Pre-ranked gene set enrichment analysis (GSEA) of mESC cell identity genes across communities ranked by signal strength for MED1, BRD4, EP300, CBP, and H3K27ac. Black tick marks indicate the community signal rank positions of genes from the “Embryonic Stem Cell” list of the PanglaoDB_Augmented_2021 database (see Supplemental Methods for additional details).

### 3D-super-enhancers

A thorough literature review uncovered that, in mESCs, condensates of specific co-activators have been shown to interact with 7 genes, all of which control cell identity, and whose associated chromatin sites are associated with SEs highly enriched for co-activator binding: *Esrrb* (MED1), *Nanog* (MED1, BRD4), *Mir290* (MED1, BRD4), *Sox2* (BRD4), *Trim28* (MED1, BRD4), *Klf4* (MED1, BRD4), and *Pou5f1* (BRD4) (4, 6, 7, 10, 72). We therefore hypothesized that the genes within CRE communities that had exceptionally high co-activator signal may be enriched for cell-identity-defining genes (4–6, 9). We began by ranking communities by their signal enrichment for MED1, BRD4, EP300, CBP, and H3K27ac. We observed an asymmetric distribution of cumulative protein signal across communities for each mark; in each case, most communities accumulated a modest amount of signal, whereas a minority of communities (<10%) accumulated an exceptional amount of signal (Fig. 2B).

Irrespective of the mark used to stratify communities, we noted qualitative similarities between how co-activator signal was distributed across communities and well-established co-activator signal distributions at CRE clusters assembled by genomic proximity (13, 14, 39). Exceptionally high-signal linear CRE clusters have been studied in great detail as “super-enhancers;” as an evolution of this concept, we chose to term the exceptionally high-co-activator subset of our topology-based CRE communities “3D super-enhancers”, or 3D-SEs (Supp. Data S6). To define this subset mathematically, we used the established signal cutoff strategy from the linear SE software, ROSE (Methods) (13, 14, 39). All but one known co-activator condensate-associated genes were in 3D-SEs defined by all analysed co-activators; the lone exception was *Trim28*, which was associated with MED1-ranked, BRD4-ranked, and H3K27ac-ranked 3D-SEs but was sub-threshold when communities were ranked by EP300 and CBP (Fig. 2B). Intriguingly, the numbers of 3D-SEs for MED1 and BRD4 (1,158 and 986) more closely comport with the number of observed MED1 and BRD4 protein foci in mESC immunofluorescence (∼1,000) than the number of MED1 or BRD4 linear SEs (214 and 422), which suggests that 3D-SEs might correspond more comprehensively to co-activator condensates (Supp Data S18, S20) (4).

3D-SEs had a greater degree of factor-specificity than moderate-signal communities (Fig. 2C). An analysis of the coefficient of variance for 3D-SEs using their relative enrichment in MED1, BRD4, EP300, and CBP showed that a subset of communities was relatively rich in certain co-activators and relatively poor in others (Supp. Fig. 3E, F, Supp. Data S21, Supplemental Methods). This observation suggests that not all genes that are associated with one type of condensate might be expected to be associated with other types of condensate.

Though both were derived by identifying signal-rich CRE collections, we noted interesting differences between linear SEs and 3D-SEs (Supp. Fig. 4A). For example, the histone gene clusters are predominantly regulated by promoter elements and are also known to form the biomolecular condensates known as the “Histone Locus Bodies” (73). However, despite their enrichment for marks of activating chromatin, these gene clusters have not been associated with linear SEs (Supp. Fig. 4B, C) (73). Consistent with its apparent chromatin signal, *H2ac10*, a representative gene from this cluster and one of 23 total histone genes associated with MED1 3D-SEs, is associated with 3D-SEs defined by all analysed factors (Fig 2C). Thus, the strategy underpinning BOUQUET enables assignment of high signal to condensate-associated genes with distinct CRE network properties.

We performed a gene ontology analysis of the genes associated with MED1 3D-SEs. As anticipated, the most significantly enriched term was “Embryonic Stem Cells.” (Fig 2D, Supp. Data S22). We next compared MED1-identified 3D-SEs with those identified by BRD4, EP300, CBP, or H3K27ac. We found that the most significantly enriched term was consistently “Embryonic Stem Cells” (Fig. 2D, Supp. Data S22). Indeed, there was a significant genome-wide correlation between community signal loading and ESC-relevant genes via pre-ranked Gene Set Enrichment Analysis, and this relationship remained consistent across the different loop-calling thresholds we examined (Fig. 2E, Supp. Fig. 4D, Supp. Data S23). Thus, 3D-SEs comprise CREs and target genes that assemble large amounts of specific co-activator apparatus and are associated with cell-identity-defining genes and other extremely highly regulated genes.

### 3D-SEs are enriched in cell identity genes across mammalian tissues

We next sought to determine whether 3D-SEs were specific to ESCs or could be found in other mammalian tissues. To enable community identification across a broad range of systems, we designed BOUQUET to be compatible with a variety of input datatypes. For use in tissues and cells where the preferred datasets may be unavailable, we implemented a minimal protocol (BOUQUET-lite) that only requires H3K27ac HiChIP, which generally reflects topological interactions, CRE activity, and gene expression, albeit with certain limits (Methods). The characteristics we observed in mESC “full” BOUQUET communities were generally consistent with communities derived from BOUQUET-lite. Specifically, both protocols uncovered comparably shaped distributions of signal across communities genome-wide, a high correlation between community signal and community gene expression, and recovery of known co-activator condensate-associated genes within 3D-SEs (Supp. Fig. 5A, 5B, Supp. Data S15). As such, we propose that BOUQUET-lite is an appropriate substitute for studying communities in systems with limited data.

To interrogate CRE communities in differentiated mammalian cell types, we executed BOUQUET-lite on published H3K27ac HiChIP datasets in mESCs and four other mouse cell/tissue types: cardiomyocytes, somatostatin (SST)-expressing neurons, thymocytes, and regulatory T cells (Treg) (Supp. Data S10-S14) (74–77). Consistent with our observations in full BOUQUET for mESCs, cumulative community H3K27ac signal was significantly positively correlated with community gene expression in all cell/tissue types (Fig. 3A). We noted asymmetric distributions of H3K27ac signal across communities genome-wide, with small fractions of communities that were exceptionally signal-rich in each cell/tissue type (Fig. 3B). Altogether, these results demonstrate that co-activator–rich CRE communities are a general phenomenon found across tissue types.

**Figure 3:**
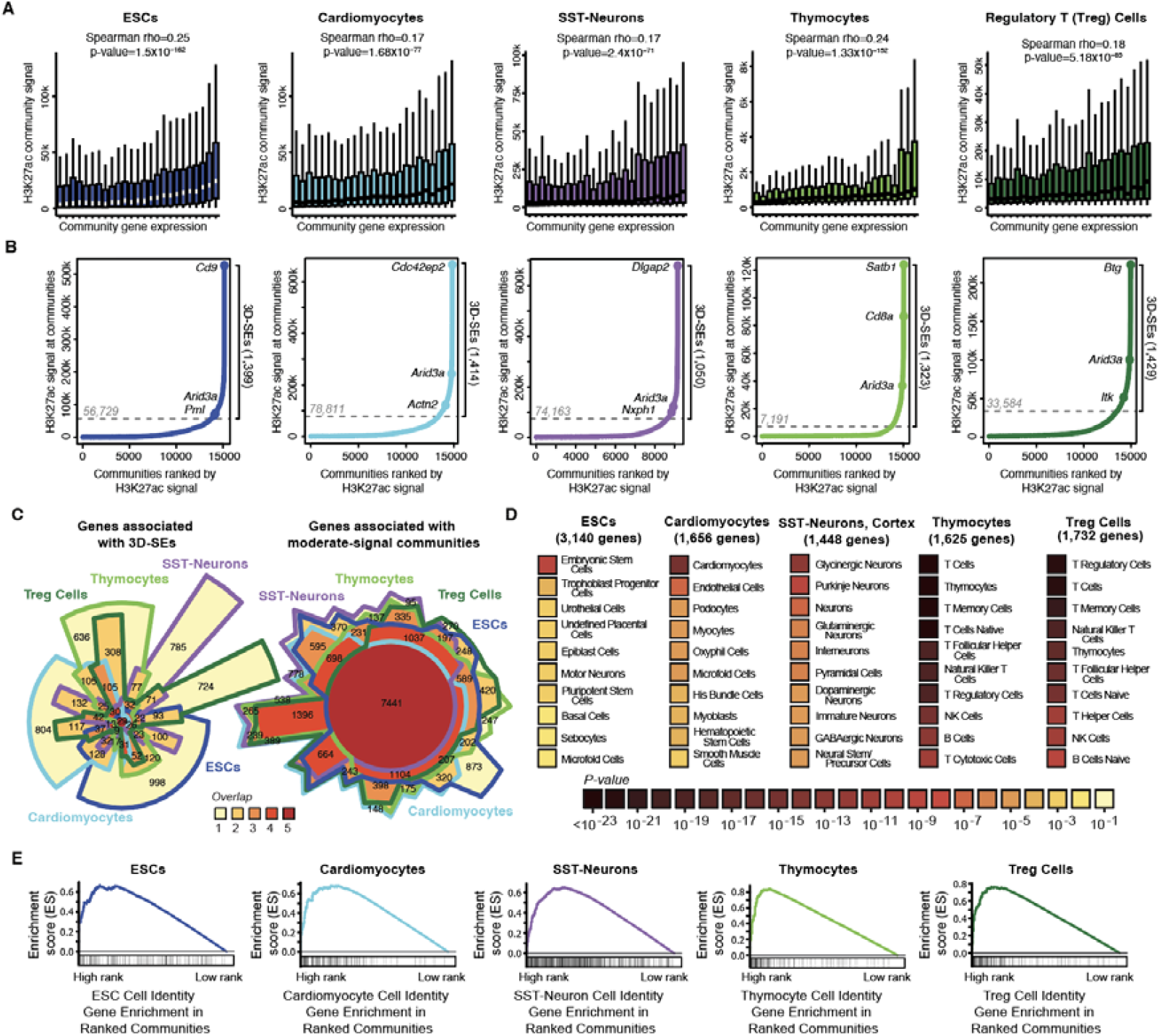
3D-SEs mark cell-identity-defining genes across tissues. (**A**) Cumulative per-community H3K27ac HiChIP signal versus focal gene expression for five mouse cell/tissue types: mouse embryonic stem cells (mESCs), cardiomyocytes, somatostatin (SST)-producing neurons, thymocytes, and regulatory T (Treg) cells. Communities are binned into 25 groups by increasing focal gene expression levels. (**B**) Distribution of H3K27ac HiChIP signal ranking across CRE communities in five mouse cell/tissue types. Y-axis: per-community signal quantified as summed H3K27ac HiChIP reads across community CREs; X-axis: relative community rank by per-community signal; Example genes are labeled, including the highest-ranked gene, a representative cell-identity-defining gene, and a gene identified as 3D-SE-associated in five of five analyzed cell/tissue types (*Arid3a*). Total count of 3D-SEs is shown on the right; dashed line marks cutoff distinguishing 3D-SEs from moderate-signal communities. A minority of communities exceed this 3D-SE threshold in each cell type: 10.2% in mESCs, 10.5% in cardiomyocytes, 7.9% in somatostatin (SST)-expressing neurons, 9.7% in thymocytes, and 10.6% in regulatory T (Treg) cells. (**C**) Chow-Ruskey diagrams showing the overlap of genes associated with moderate-signal communities (left) or 3D-SEs (right) in combinations of five cell/tissue types. The area of each section of the diagram is proportional to the number of genes within it. (**D**) Gene ontology analysis for ‘cell type’ term enrichment in 3D-SE genes across five cell/tissue types. The top 10 most enriched terms from the PanglaoDB_Augmented_2021 database are shown for each cell/tissue type (see Supplemental Methods for additional details). (**E**) Pre-ranked gene set enrichment analysis (GSEA) of cell identity genes across five mouse cell/tissue types. Black tick marks indicate the community signal rank positions of genes from significantly enriched cell identity terms in the PanglaoDB_Augmented_2021 database (see Supplemental Methods for additional details).

We next examined the genes associated with CRE communities and their specificity across cell types. Genes associated with 3D-SEs tended to be associated with 3D-SEs in only one cell type, whereas genes in moderate-signal communities were largely shared across multiple cell/tissue types (Fig. 3C, Supp. Data S15). Consistent with this pattern, heatmap analysis of publicly available RNA-seq demonstrated cell-type-specific expression of 3D-SE-associated genes (Supp. Fig. 5C), with elevated expression specifically in their corresponding cell type (except for the closely related Thymocyte and Treg cell types). Though certain genes were in 3D-SEs in multiple cell types, including several groupings of histone genes, we observed variability in the structures of those 3D-SEs (Fig. 3B, Supp. Fig. 6A-C, Supp. Data S15). This observation suggests that genes are generally associated with 3D-SEs in a cell-type-specific manner and that, when associated with 3D-SEs in multiple cell types, their CRE composition can also be cell-type-specific.

If genes are associated with 3D-SEs in a cell-type-specific manner, it would be expected that these genes have roles in executing or establishing those cell identities. We performed gene ontology analyses in each cell/tissue type to determine whether 3D-SE-associated genes are enriched for genes associated with identity-defining functions across the body (Fig. 3D, Supp. Data S22) (4–7, 9). Genes associated with 3D-SEs were indeed specifically enriched for signature factors known to contribute to specialized cell identities and functions, consistent with their source cell/tissue type, e.g., *Pml* in mESC, *Actn2* in cardiomyocytes, *Nxph1* in SST-neurons, *Cd8a* in thymocytes, and *Itk* in Treg cells (Fig. 3B, Supp. Fig. 7A, 7B) (78–82). Further, the composition of these 3D-SEs, including both constituent CREs and the connections among them, were cell-type-specific. Pre-ranked Gene Set Enrichment Analysis showed that, in a given cell type, elevated community signal was significantly linked to roles in cell identity (Supp. Data S23) (Fig. 3E). In mESCs, we noted that certain housekeeping genes, including those encoding histones, were also associated with 3D-SEs. We therefore hypothesized that genes with multi-tissue 3D-SE associations might exhibit enrichment for housekeeping functions. We tested this hypothesis using a curated list of 1,128 housekeeping genes from MSigDB and found a significant enrichment relative to expectation (P = 9.6 × 10^-12^, hypergeometric test) (70). Together, these results suggest that the regulatory communities associated with high levels of transcriptional machinery are largely enriched for cell identity genes, as well as highly expressed genes in gene-rich regions that encode for housekeeping factors, across mammalian tissues.

### 3D-SEs inform expression responses to signaling pathways

The heterogenous binding of TFs at CREs has been proposed as a mechanism by which a variety of cellular signals can be integrated into a gene’s expression, especially for key cell identity genes associated with multiple CREs (63). Moreover, co-activator condensates have been observed to sequester high concentrations of signaling transcription factors (sTFs) (72). We thus hypothesized that 3D-SEs integrate cellular signals and enable elevated community-spanning responses to multiple signaling pathways. To test this hypothesis, we reanalysed ChIP-seq data from three terminal TFs of notable signaling pathways relevant in mESCs: STAT3, SMAD3, and TCF3 (83–85). We examined the CREs of 3D-SEs, moderate-MED1-signal communities, and size-matched randomized control sets of CREs, to determine whether each set of CREs was collectively bound by 0, 1, 2, or 3 types of signaling TFs (Supplemental Methods; Supp. Fig. 8A). Most 3D-SEs (60%) contained binding events for all three types of sTF, which was significantly more than moderate-MED1-signal CRE communities (8%) or randomized communities of the same size (39%, two-sided Chi-squared test P-value=1.5×10^-49^). Further, 3D-SE genes showed a greater change in gene expression following signaling pathway perturbation: TGF-β pathway perturbation (suppressing expression by blocking Tgf-β with SB431542; p = 2.9×10^-15^), LIF pathway stimulation (activating expression by exposing to LIF for 1h; p = 3.9×10^-7^), and Wnt pathway perturbation (activating expression by *Tcf3* knockdown; p=5.4 ×10^-9^) (Supp. Fig. 8B, Supp. Data S24). Interestingly, 3D-SE genes were more dramatically responsive to signaling perturbation than the genes that were the nearest linear neighbors of signaling-TF-bound CREs (Supp. Fig. 8C, Supp. Data S23). Altogether these results demonstrate that 3D-SEs amplify the gene expression responses to critical transcriptional signaling pathways by accumulating heterogeneous CREs.

### 3D-SEs as transcriptional condensate components

Several transcriptional proteins, including MED1 and BRD4, form high-density condensates that are observable in fluorescence microscopy experiments as punctate nuclear signals within a dilute background (2, 4, 6, 7). Based on their exceptional protein concentrations, we hypothesized that the CREs of 3D-SEs would interact with co-activator puncta, thus establishing BOUQUET as a genome-wide identifier of condensate-associated genomic regions. To test this hypothesis, we used fluorescence microscopy to study the largest 3D-SE in mESCs, ranked 1^st^ by MED1 signal, and 3^rd^ by BRD4 signal (Fig. 4A, Supp. Fig. 9A). This community comprises three Insulated Neighborhoods that each contain at least one signature protein-coding gene, *Esrrb, Irf2bpl*, and *Sptlc2*, so we refer to it as the EIS community. The EIS community spans ∼1MB, which minimizes the likelihood that co-localization observations between elements at its most extreme ends would arise from stochastic chromatin movements. The source chromosome of this community is chr12, which has two alleles in our mESCs (Supp. Fig. 9B). The EIS community locus is therefore ideal to study the interactions and activities of a broad, protein-rich community.

**Figure 4:**
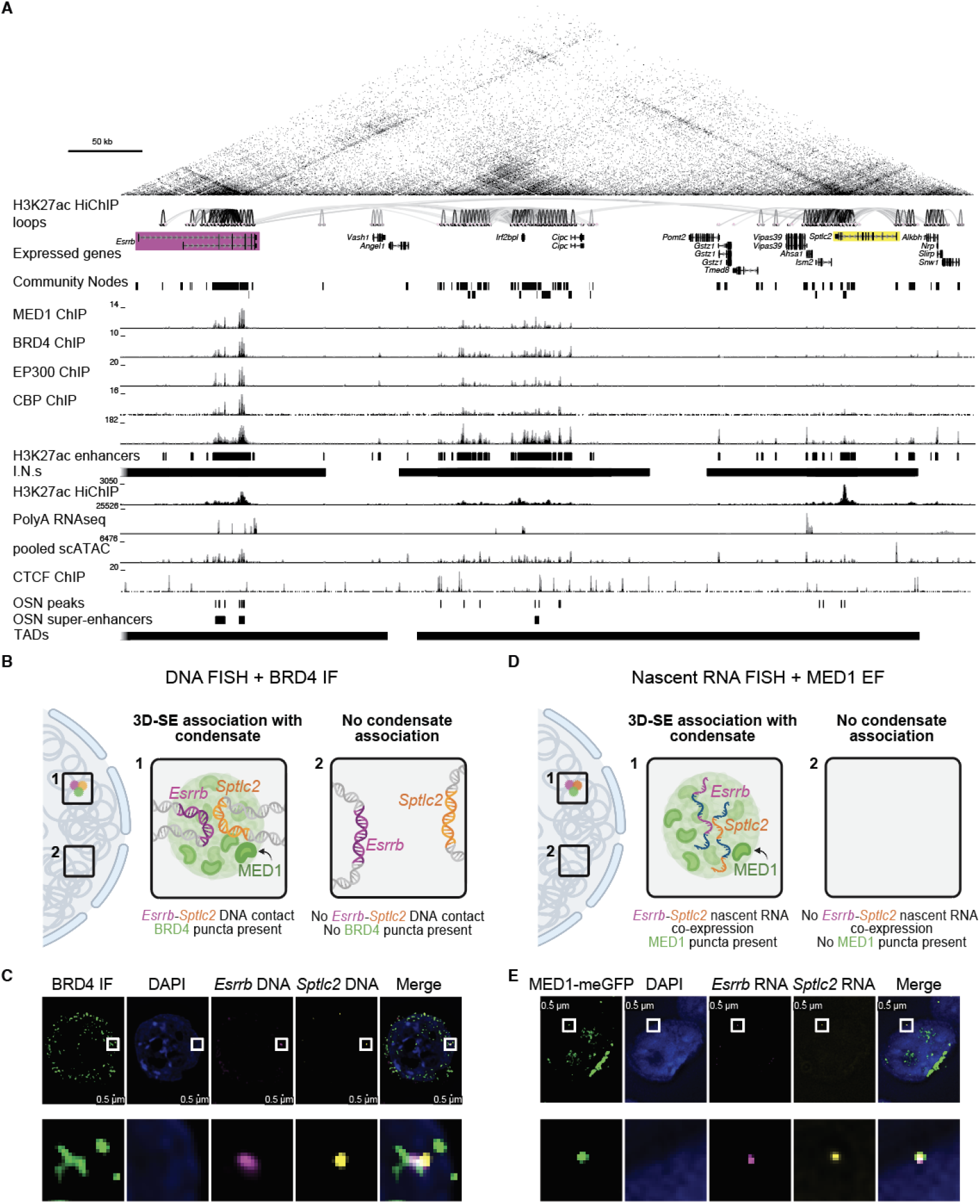
3D-SEs colocalize with putative co-activator condensates. (**A**) UCSC Genome Browser view of the locus containing the *Esrrb/Irf2bpl/Sptlc2* 3D-SE, the target of the DNA and RNA FISH experiments. ChIP-seq tracks are reads-per-million (RPM) normalized. (**B**) Model of two predicted outcomes for the DNA FISH and BRD4 immunofluorescence (IF) experiments. In case 1 (*box 1, middle*), the DNAs of two genes are associated within a BRD4 co-activator punctum (*nuclear diagram, left, region of interest (ROI) 1*). In case 2 (*box 2, right*), the DNA are not associated in the ROI (*nuclear diagram, left, ROI 2*). In the model and below, *Esrrb, Sptlc2*, and BRD4 are represented in magenta, yellow and green respectively. Created in BioRender. Maher, K. (2025) https://BioRender.com/fghhosi. C) Fluorescence microscopy of mESC nuclei following DNA FISH with BRD4 IF. Blue = DAPI stain for chromatin; magenta = DNA probes for *Esrrb*; yellow = DNA probes for *Sptlc2*; green = anti-BRD4 antibody. Images represent a single z-plane. Scale bar for top panels = 0.5 *μ*m. Lower panels are zoomed-in views of the same ROI shown in the top panels. (D) Model of two predicted outcomes for the RNA FISH and endogenously tagged MED1 fluorescence (EF) experiments. In case 1 (*box 1, center*), the mRNAs the two genes encode are associated within a MED1 co-activator punctum (*nuclear diagram, left, ROI 1*). In case 2 (*box 2, right*), the mRNAs are not co-expressed nor associated with MED1 punctum (*nuclear diagram, left, ROI 2*). In the model, *Esrrb, Sptlc2*, and BRD4 are represented in magenta, yellow and green respectively. Created in BioRender. Maher, K. (2025) https://BioRender.com/ycwql9z. (E) Fluorescence microscopy of mESC nuclei following RNA FISH with MED1 EF. Blue = DAPI stain for chromatin; magenta = nascent RNA probes for *Esrrb*; yellow = nascent RNA probes for *Sptlc2*; green = MED1 EF. Images represent a single z-plane. Scale bar for top panels = 0.5 *μ*m. Lower panels are zoomed-in views of the same ROI shown in the top panels.

All genes that have been observed to interact with co-activator condensates had been previously assigned to linear super-enhancers, e.g., *Esrrb* from the EIS community. However, the closest linear super-enhancer to *Sptlc2* is over 400 kb away, and it is instead assigned to *Irf2bpl* (Supp. Data S18). To determine if the high community signal associated with *Sptlc2* by BOUQUET accurately predicted the interactions of *Sptlc2* with co-activator condensates, we performed DNA FISH and anti-BRD4 immunofluorescence in >200 mESCs (Fig. 4B, Supp. Fig. 9A). We searched 0.4 μm regions of interest (ROIs) centered on *Sptlc2* gene signal and found 6.9% (26 out of 376) indeed co-localized in this small region with punctate BRD4 co-activator signal (Fig. 4C, Supp. Fig. 9C). Unlike *Sptlc2, Esrrb* has been assigned a linear SE, which is intragenic to *Esrrb* (Supp. Data S18). As a comparison, the frequency of co-localization between *Esrrb* and BRD4 puncta was modest (2.7%, 11/401) (Supp. Fig. 9C), despite previous deep study of the interactions between *Esrrb* and co-activator puncta (6). Thus, BOUQUET can identify 3D-SEs that predict the interaction of unexpected genes with co-activator condensates.

Since *Esrrb* and *Sptlc2* are within the same 3D-SE and have been shown to interact with co-activator puncta, we hypothesized that both genes could interact with the same individual punctum (4, 6). 0.4 μm ROIs were constructed around *Sptlc2* or *Esrrb* DNA FISH puncta, and we characterized their co-localization as well as their tripartite interactions with BRD4 puncta. The genes that encode *Esrrb* and *Sptlc2* frequently co-localized in these small regions, and several of these co-localized occurrences also overlapped BRD4 foci (Fig. 4C, Supp. Fig. 9C). As a negative control for our microscopy-based co-localization analysis, we performed DNA FISH for *Tmed10* in our mESCs. *Tmed10* is as highly and frequently expressed as *Esrrb* and *Sptlc2* are; it is approximately the same distance from *Esrrb* as *Sptlc2* (Supp. Fig. 9D-F). *Tmed10* and *Esrrb* are not connected by high-confidence loops, nor do they appear in the same BOUQUET community, though there are observed contacts between *Tmed10* and *Esrrb* (Supp. Fig. 9D-F). In single cells, *Esrrb* co-localized ∼20% less frequently with *Tmed10* (24, 5.6% of *Esrrb* ROIs) than with *Sptlc2* (29, 7.2% of *Esrrb* ROIs). Furthermore, *Tmed10*/BRD4 co-localization (0.29%, 1/348 *Tmed10* ROIs overlapping BRD4) was substantially lower than *Esrrb*/BRD4 co-localization (1.9%, 8/426 *Esrrb* ROIs overlapping BRD4) (Supp. Fig. 9C,G). Tripartite interactions of *Sptlc2*/*Esrrb*/BRD4 were observed 10 times (1.3% of 748 gene ROIs). In contrast, 0 tripartite interactions of *Tmed10*/*Esrrb*/*BRD4* were observed (0 out of 750 ROIs). Together, these results support a model in which *Sptlc2* and *Esrrb* interact simultaneously within the same individual co-activator punctum in a non-random manner.

Using higher-resolution experiments, we next examined how co-localization of genes related to coordinated transcriptional regulation. Single-cell RNA-seq demonstrated that most cells contained transcripts of both *Esrrb* and *Sptlc2* (77%), and single-cell ATAC-seq demonstrated that both of their promoters were transposase-accessible (64%) in the vast majority of cells (Supp. Fig. 9F). However, these assays do not directly interrogate the coordination of gene transcription or CRE activity, nor how this coordination relates to co-activator signal. To measure coordinated transcription, we designed RNA FISH probes to mark only the nascently transcribed RNAs of *Esrrb* and *Sptlc2* and studied their expression patterns in mESCs bearing endogenously tagged meGFP-MED1 (Fig. 4D, Supp. Fig. 9B,H, Supplemental Methods). These RNA probes therefore also mark the precise nuclear region containing the gene being transcribed.

We defined 627 regions of interest (ROIs) as 0.4 μm nuclear regions that contained a punctum for at least one nascent transcript. Most co-activator puncta and their associations with genes are short-lived, and RNA transcription is infrequent and bursty, complicating analysis of their coincidence (6, 9, 86). To establish a background expectation, we examined the 291 ROIs that were positive for *Esrrb* transcription and observed that only 4.1% of such ROIs coincided with punctate MED1 signal (Fig. 4E, Supp. Fig. 9I). We observed a comparable fraction, 3.6% of 365 ROIs with *Sptlc2* transcription, coincided with punctate MED1 signal. Thus, *Sptlc2* interacts with co-activator puncta with comparable frequency as *Esrrb* whose interactions with co-activator puncta have been detailed.

We next examined the relationship between co-transcription and co-activator puncta. As expected, only 4.6% of 627 ROIs contained both *Esrrb* and *Sptlc2* transcription. Consistent with the observation that *Esrrb* and *Sptlc2* DNA are likely to interact within the same BRD4 punctum, we see that in ROIs with co-transcription, 17% (5/29) showed RNA co-localization with MED1 puncta (Fig. 4E). We repeated this experiment with cells bearing BRD4-meGFP and also observed three-way co-localization between nascent *Esrrb* RNA, nascent *Sptlc2* RNA, and BRD4 puncta in these cells (Supp. Fig. 9J). Thus, 3D-SEs contain genes that are co-transcribed and co-localize with the same co-activator punctum. Altogether, via genome-wide unbiased analysis, BOUQUET identifies genes that interact with co-activator puncta at similar frequency as genes previously studied in great detail for their interactivity with such puncta.

## DISCUSSION

Understanding which cis-regulatory elements (CREs) and associated transcriptional machinery regulate each gene is critical to decoding gene regulation and cell identity. Popularly used methods for unbiased assignment of large collections of regulatory apparatus to target genes and CREs, such as ROSE and LILY, are limited in their ability to incorporate chromatin topology. Furthermore, these approaches neither capture the full complexity of CRE function nor fully account for the transcriptional apparatus relevant for a given gene’s regulation. For instance, promoter exclusion done by ROSE as a default precludes discovery/incorporation of the known phenomenon wherein promoters act as enhancers of other genes (34). Attempts to predict the regulatory contributions of given CREs by combining individual CRE activity and their direct genomic contacts have uncovered several quantitative relationships that connect chromatin measurements like contact frequency and protein-binding to expression, but these approaches ultimately do not interrogate the known multi-faceted interactions of multiple enhancers with multiple promoters observed in mammalian cells (64). A more comprehensive approach would capture the entire set of regulatory elements that impact a given gene’s expression, irrespective of their proximity in a linear genome or stratification between enhancer and promoter. This philosophy has guided other groups to develop models of transcriptional “hubs” or “highly interconnected enhancers” that represent individual CREs with many interaction partners (16, 17, 38). While the hub model can identify sets of CREs associated with a single focal node using direct loop connections, recent implementations enforce linear contiguity of hubs, thus missing subdivisions within these stretches, broader context of how genes and CREs throughout that same region interact with each other in interconnected networks, and how these arrangements result in spatiotemporal organization of protein apparatus controlling transcription. Further, these approaches are ultimately limited by statistical challenges in setting appropriate significance thresholds for loop-identification, since they exclude connections that are sub-threshold. The limitations of these approaches inspired us to develop a strategy to optimally assemble CREs and associated protein apparatus.

Here, we describe BOUQUET, a novel computational framework that quantifies the transcriptional protein apparatus associated with a given gene by detecting the gene’s regulatory community via well-characterized categories of topological interaction. BOUQUET therefore incorporates measurements of genome organization and transcriptional regulation, namely network-based community detection for cutoff-free construction of CRE sets, which reduces reliance on error-prone and sometimes non-representative loop calls used to identify HICEs and hubs, all of which supersedes the linear genomic arrangement assumption underpinning tools for protein assignment like ROSE.

Recent studies have revealed that collections of co-activator protein within the nucleus often are associated with liquid-liquid phase-separated (LLPS) droplets or transcriptional condensates; additional physical models and terms have been proposed and are under ongoing study. Irrespective of biophysical process, being able to predict which genes and regulatory elements may functionally associate with these protein foci is of prime interest. The field is limited in its studies because fewer than 10 genes have been observed to associate with co-activator condensates, but more than a thousand co-activator foci have been observed per nucleus. Other chromatin proteins that form condensates may have similar discrepancies. To our knowledge, BOUQUET represents the first tool to infer genome-wide CRE collections and link them systematically to transcriptional condensates. To our surprise, BOUQUET uncovered a significant correlation between the amount of protein apparatus we could assign to a gene’s community and that gene’s expression; this relationship is intuitive but (to our knowledge) has not been reported, which we suppose is due to long-standing difficulties with CRE-gene assignment. The uneven or asymmetric distribution of proteins at communities enabled us to identify an extremely protein-rich subset of communities we refer to as “3D-super-enhancers.” 3D-SE genes include all known condensate-associated genes, and the number of 3D-SEs roughly matches the number of condensates. We demonstrate that at least one gene not previously associated with a linear SE is condensate-associated, suggesting that 3D-SEs might correspond to co-activator condensates.

Unsurprisingly, many of our 3D-SE genes included many of the same cell-identity-defining genes previously identified as associated with linear SEs; however, through the inclusion of topology, we can more expansively characterize the gene regulatory interactions associated with those genes and nominate additional CREs putatively involved in regulating their expression. It is likely that each CRE within a community contributes to the regulation of each gene within that same community in a distinct manner, including the possibility of redundancy, additivity, synergy, competition, and/or hierarchy among community CREs, especially as they respond to distinct sets of signaling pathways (76, 87). Here, we proposed additivity of signal among activating CREs would be a reasonable measure of community activity, but other strategies to combine CREs of different types and their associated signal may refine our findings. We are also able to identify high-signal communities overlooked by enhancer-biased alternatives, such as the promoter-heavy condensate-forming histone gene cluster (73). Based on our findings, it also seems likely that multiple SE-associated genes can be co-regulated within the context of a single community, such as the *Esrrb-Irf2bpl-Sptlc2* 3D-SE community (Supp. Data S5, S18). This finding suggests that the study of mutant enhancers should similarly expand if topology is considered and should improve the prediction of the effects of non-coding mutation.

It has been observed that co-localized genes are co-expressed and that co-localized enhancers are coordinately active (88, 89). Our CRE communities provide an effective framework for identifying CREs whose regulatory activities are linked. Indeed, we observed a colocalization between co-activator puncta and multiple components of the largest 3D-SE (containing *Esrrb; Irf2bpl; Sptlc2;* et al.) when multiple constituent genes were both expressed and in proximity. These observations demonstrate that 3D-SEs can be observed in single cells, are not merely an artifact introduced by HiC technologies, and that multiple genes can be arranged near or in the same co-activator punctum. Our microscopy data suggest that CREs collaborate to regulate gene expression within a 3D-SE through multi-partite physical interactions. Several technical challenges limit our assessment of co-localization with immunofluorescent imaging, including probe intensity, photobleaching, relocation misalignment, and the stringency of our analysis using sub-micron regions of interest. This latter concern is reflected by the field’s inconsistent reporting of frequencies of interactions and range of distance cutoffs for elements being “co-localized.” We therefore believe our evaluations underestimate the close and frequent spatial arrangements of co-activator puncta and CRE communities. Further, transcriptionally productive associations between condensates and their gene targets have been shown to be highly transient (∼12 seconds), increasing the likelihood that co-localization events may be too brief to be captured by static imaging approaches (6). Nonetheless, the findings of this study provide a valuable direct comparison of the co-localization of nucleic acids as measured by two commonly used orthogonal approaches, microscopy and sequencing-based chromosome conformation capture. Future efforts will be needed to disentangle the roles of different CREs and regulatory proteins in governing gene expression within these complex networks, as well as the kinetics and frequency of each event, including punctum lifecycles. Ultimately, by accounting for the various layers of physical arrangement of the nucleus—chromatin composition, genome structure, and condensate assembly—when tallying the transcriptional apparatus that is relevant to the regulation of each gene, we can more accurately reflect biological reality and offer new insight into the fundamentals of gene regulation.

## Supporting information

Supplemental Combined

## ACKNOWLEDGEMENTS

We thank Richard A. Young (Whitehead Institute) for providing parental mESC V6.5 and deletion (dMir290e) mESC V6.5 lines. We thank Charles H. Li, Abraham S. Weintraub, and Tong Ihn Lee for early conceptual discussions. We thank Sarah August for the editing support.

## AUTHOR CONTRIBUTIONS

Jie Lv: Conceptualization, Data Curation, Formal Analysis, Investigation, Methodology, Software, Visualization, Writing – Original Draft. Kelsey A. Maher: Data Curation, Formal Analysis, Investigation, Software, Visualization, Writing – Original Draft. Li Dong: Investigation. Virginia Valentine: Investigation. Seth Staller: Investigation, Software. Alaguraj Veluchamy: Data Curation, Formal Analysis, Software. Li Tian: Investigation. Yuna Kim: Investigation. Bensheng Ju: Investigation. Marcus Valentine: Resources, Visualization. John Easton: Resources. Stanley B. Pounds: Formal Analysis, Methodology, Resources, Software, Visualization. Steven Burden: Formal Analysis, Investigation, Resources, Visualization. Brian J. Abraham: Conceptualization, Formal Analysis, Funding Acquisition, Methodology, Software, Supervision, Visualization, Writing – Original Draft.

## SUPPLEMENTARY DATA

Supplementary Data are available at *NAR* Online.

## DATA AVAILABILITY

Data generated in this study have been deposited to the Gene Expression Omnibus (GEO) under accession number GSE254728. Publicly available datasets used in this study are detailed in Supplemental Data S28. UCSC Genome Browser tracks can be viewed at: https://genome.ucsc.edu/s/abraham_lab/3DSE_mm10. Code generated in this study is available at the following Github repositories: Hi-LOW (https://github.com/stjude/HiLOW; 10.5281/zenodo.15786041); BOUQUET (https://github.com/stjude/BOUQUET/; 10.5281/zenodo.15782836); LASER (https://github.com/stjude/LASER; 10.5281/zenodo.15786043); Insulated Neighborhood Caller (https://github.com/stjude/Dowen_Fan; 10.5281/zenodo.15784157).

