## Supplemental Combined for "3D-super-enhancers are condensate-associated cis-regulatory communities"

Supplemental Figure 1

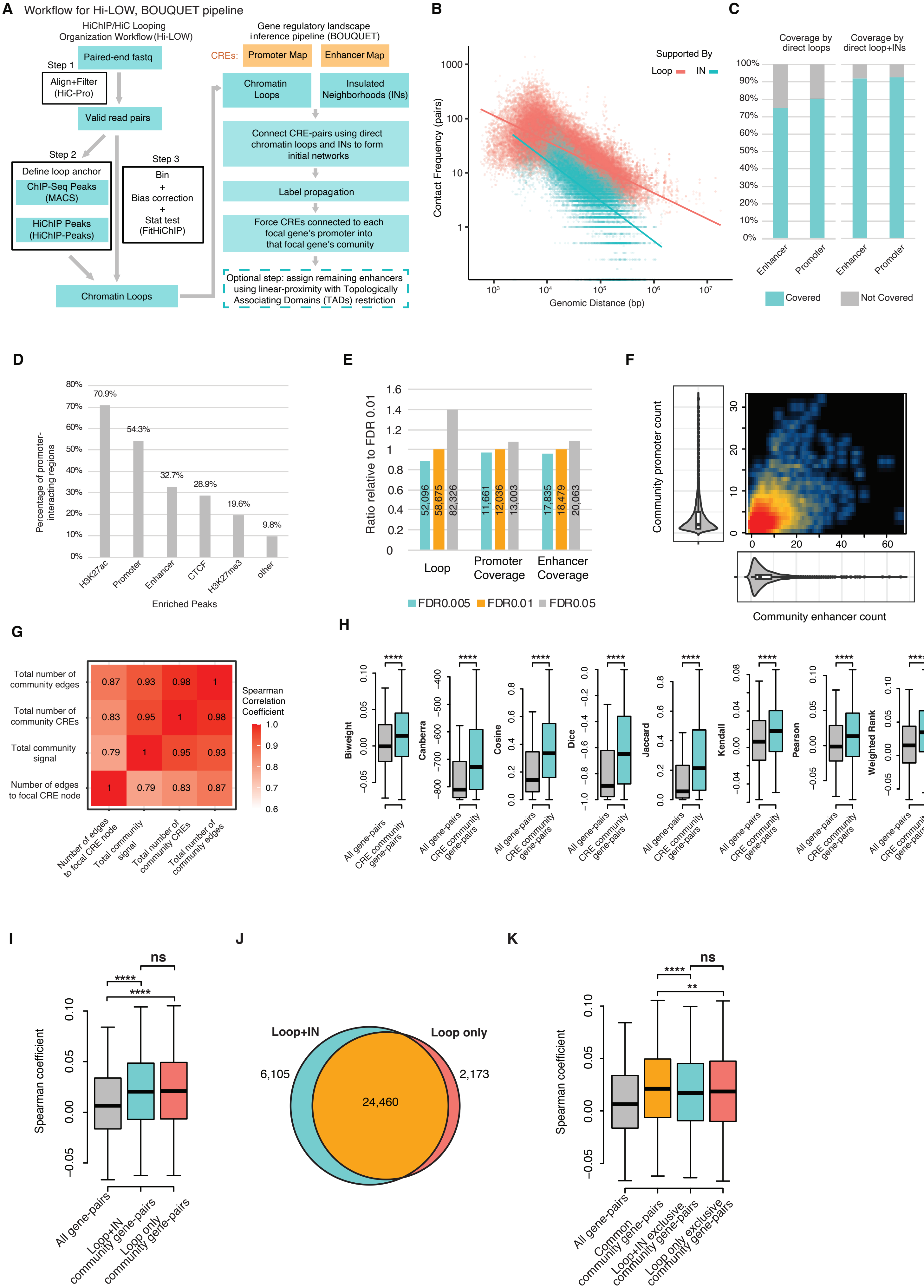

**Figure S1: BOUQUET combines protein loading with genome topology to model cis-regulatory communities.** (**A**) Computational workflows of HiChIP/HiC Looping Organization Workflow (Hi-LOW), which simplifies HiChIP loop-calling, and Building Optimized Units of QUantified Enhancer Topologies (BOUQUET), which identifies communities from cis-regulatory element (CRE) networks. Briefly, Hi-LOW ingests paired-end FASTQ files containing chromatin contact data, here in the form of H3K27ac HiChIP data. Utilizing the computational tools HiC-Pro, MACS, and HiChIP-Peaks, Hi-LOW generates high-confidence chromatin loop calls. BOUQUET ingests these chromatin loops and lists of promoters, enhancers, and insulated neighborhoods (INs) to connect pairs of CREs and draw genome-wide CRE networks. The initial CRE network is refined using label-propagation to generate a final CRE community for each gene. See Methods for details. (**B**) Scatterplot comparing the contact frequencies of CRE pairs connected by high-confidence H3K27ac HiChIP loops (pink) or INs (blue) plotted against the genomic distance separating the paired CREs. CRE-pairs whose connection is supported by both loops and INs are classified as "Loop-supported" (pink). (**C**) Fractions of enhancers and promoters that are covered by direct H3K27ac HiChIP loops or are covered by both direct HiChIP loops and fall within INs. Blue indicates the percentage covered by the given feature; gray indicates the percentage not covered. (**D**) Characteristics of the genomic regions (5 kb bins) that interact with promoter regions by high-confidence (FDR<0.01) H3K27ac HiChIP loops. Individual promoter-interacting regions may exhibit multiple characteristics due to overlapping enriched peaks of different chromatin-binding factors. (**E**) Comparison of the numbers of loops and the numbers of promoters and enhancers involved in loops across three loop-calling FDR thresholds: 0.005, 0.01, and 0.05. Bar plots show values expressed as ratios relative to those obtained at an FDR threshold of 0.01. (**F**) Number of promoters (y-axis) versus the number of enhancers (x-axis) in each CRE community (Spearman rho = 0.46, p-value = $1.6x10^{-245}$). (**G**) Heatmap displaying pairwise Spearman correlation coefficients quantifying the similarity among CRE community attributes. See Methods for further community terminology definitions. (**H**) Distribution of gene-pair expression correlations using scRNA-seq generated from 847 mESCs across eight distinct correlation metrics. Gray: genome-wide control using all gene-pairs; blue: gene-pairs within the same CRE community. **** indicates a p-value < 0.0001. Biweight: p-value = $5.99x10^{-261}$; Canberra: p-value < $2.0x10^{-300}$; Cosine: p-value < $2.0x10^{-300}$; Dice: p-value < $2.0x10^{-300}$; Jaccard: p-value < $2.0x10^{-300}$; Kendall: p-value < $2.0x10^{-300}$; Pearson: p-value = $4.12x10^{-269}$; Weighted rank: p-value < $2.0x10^{-300}$ (see Supplemental Methods for additional details). (**I**) Distributions of scRNA-seq-derived expression correlations for gene-pairs in three overlapping categories: all pairs of genes genome-wide as a control (gray, median r= 0.0065), gene-pairs from communities derived from loop+IN (blue, median r=0.02), and gene-pairs from communities derived from loops alone (pink, median r=0.021). Whiskers represent interquartile range (IQR). Statistical significance was assessed using Wilcoxon rank-sum test, **** denotes p-value < 0.0001, and "ns" indicates not significant. (grey vs. blue, p-value < $2.0x10^{-300}$; grey vs. pink, p-value < $2.0x10^{-300}$; blue vs. pink, p-value = 0.09). (**J**) Area-proportional Venn diagram showing the overlap of gene-pairs from communities derived from Loop+IN and communities derived using loops alone. (**K**) Distributions of scRNA-seq-derived expression correlations for gene-pairs in four categories: all gene-pairs genome-wide as control (gray), community gene pairs unique to loop+IN approach (blue), community gene pairs unique to loop only approach (pink), and community gene-pairs shared by both approaches (“common”, orange). Statistical significance was assessed using Wilcoxon rank-sum test, **** denotes p-value < 0.0001, ** denotes p-value < 0.01, and “ns” indicate not significant (orange vs. blue, p-value = $2.2 \times 10^{-11}$; orange vs. pink, p-value = $7.7 \times 10^{-3}$; blue vs. pink p-value = 0.16).

Supplemental Figure 2

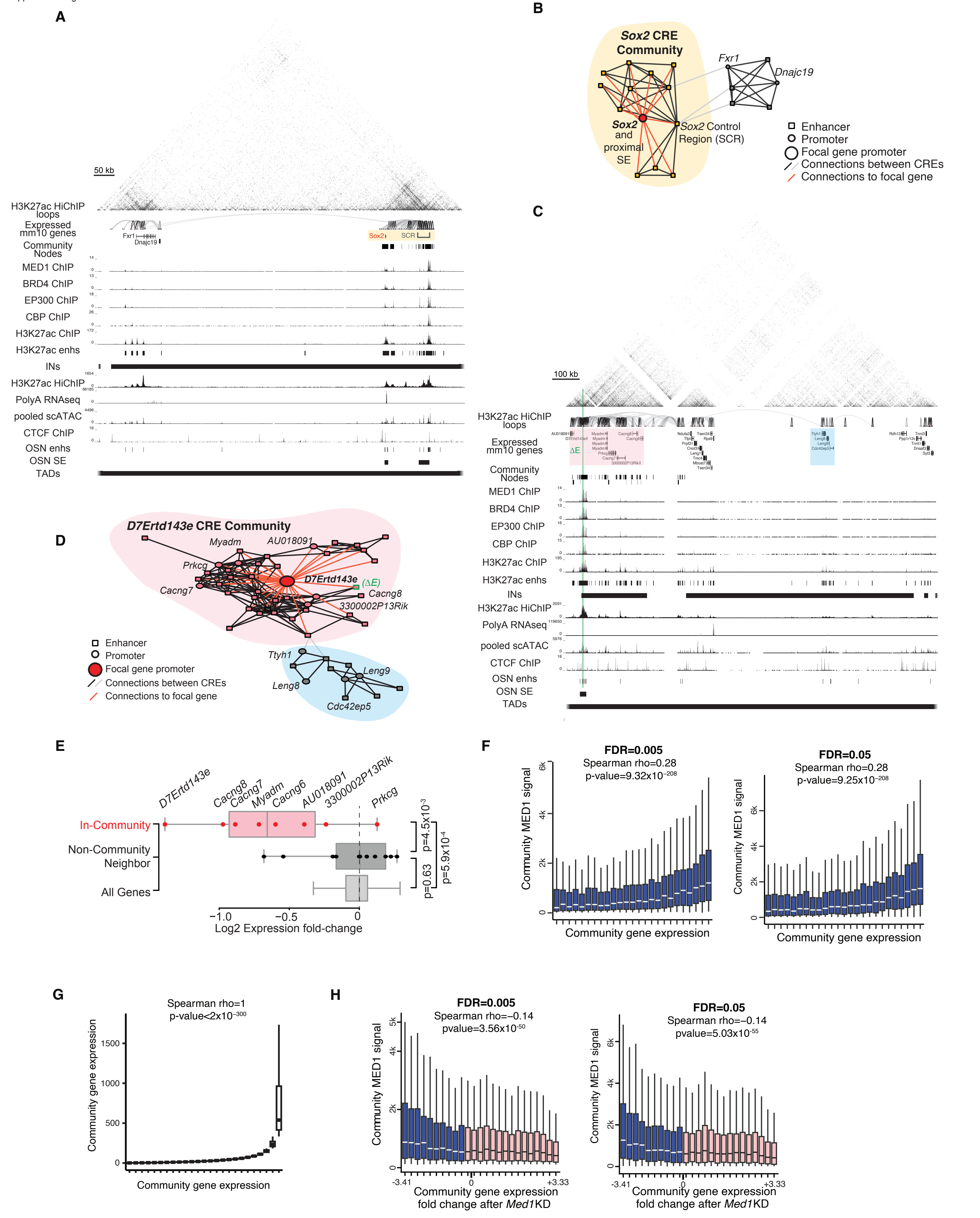

**Figure S2: CRE communities accurately model regulation of genes that interact with co-activator condensates.** (**A**) UCSC Genome Browser view of the *Sox2* component. The yellow box marks the *Sox2* CRE community. Several high-confidence H3K27ac HiChIP loops connect the promoter of *Sox2* and its proximal super-enhancer (SE) to the *Sox2* Control Region (SCR). Window size: 928 kb. ChIP-seq tracks are RPM normalized. The ends of TADs and insulated neighborhoods are faded to show that these features extend beyond the boundaries of this region. OSN SE = linear super-enhancers defined using OCT4/SOX2/NANOG as in (Whyte et *al.*, *Cell*, 2013). (**B**) Network diagram of the *Sox2* CRE component. The *Sox2* community is highlighted in yellow. While *Fxr1* and *Dnajc19* both have a single high-confidence connection to CREs of *Sox2*'s CRE community, they lack the degree of interconnection required by label-propagation to be considered as part of *Sox2*'s community. (**C**) UCSC Genome Browser view of the locus containing the *D7Ertd143e* CRE community. Genes within the *D7Ertd143e* community are marked by the red box; neighboring genes from the same component but in different communities are marked by the blue box. The CRISPR-Cas9 deletion site (ΔE) of E4, a constituent enhancer of the super-enhancer, is marked in green. Window size: 1.4 MB. ChIP-seq tracks are RPM normalized. (**D**) Network diagram of the *D7Ertd143e* component. The red cloud marks the CRE community of *D7Ertd143e*. The blue cloud marks linearly neighboring CREs that are in the same component as *D7Ertd143e*, but not within its community. (**E**) Expression changes of genes within the *D7Ertd143e* CRE community (red), neighboring genes not within the *D7Ertd143e* CRE community (dark gray), and all genes (light gray), following the deletion of the *D7Ertd143e* constituent enhancer E4. P-values of expression fold-change differences were calculated using a two-sided Wilcoxon Rank Sum Test. *D7Ertd143e*: $p=2.77\times10^{-107}$; *Myadm*: $p=6.04\times10^{-146}$; *Cacng7*: $p=2.09\times10^{-14}$; *Cacng8*: $p=3.10\times10^{-11}$; *AU018091*: $p=4.68\times10^{-45}$; *Cacng6*: $p=6.47\times10^{-4}$. **(F)** Distributions of community-assigned per-gene MED1 signal across grouped genes using alternative loop calling thresholds (FDR=0.005 left, FDR=0.05 right). Genes are sorted into 25 groups by increasing expression level (compare to Fig. 1F with FDR=0.01). **(G)** A curve representing the idealized correlation of gene expression with itself, provided for comparison with the empirical relationships shown in Fig. 1F-G and Supp. Fig. 2F). **(H)** Distributions of community-assigned per-gene MED1 signal across grouped genes following knockdown of *Med1* using alternative loop calling thresholds (FDR=0.005 left, FDR=0.05 right). Genes are sorted into 25 groups by their expression fold-change after *Med1* knock-down. Blue indicates downregulation of community gene expression following *Med1* KD; pink indicates upregulation of community gene expression following *Med1* KD (compare to Fig. 1H with FDR=0.01).

Supplemental Figure 3

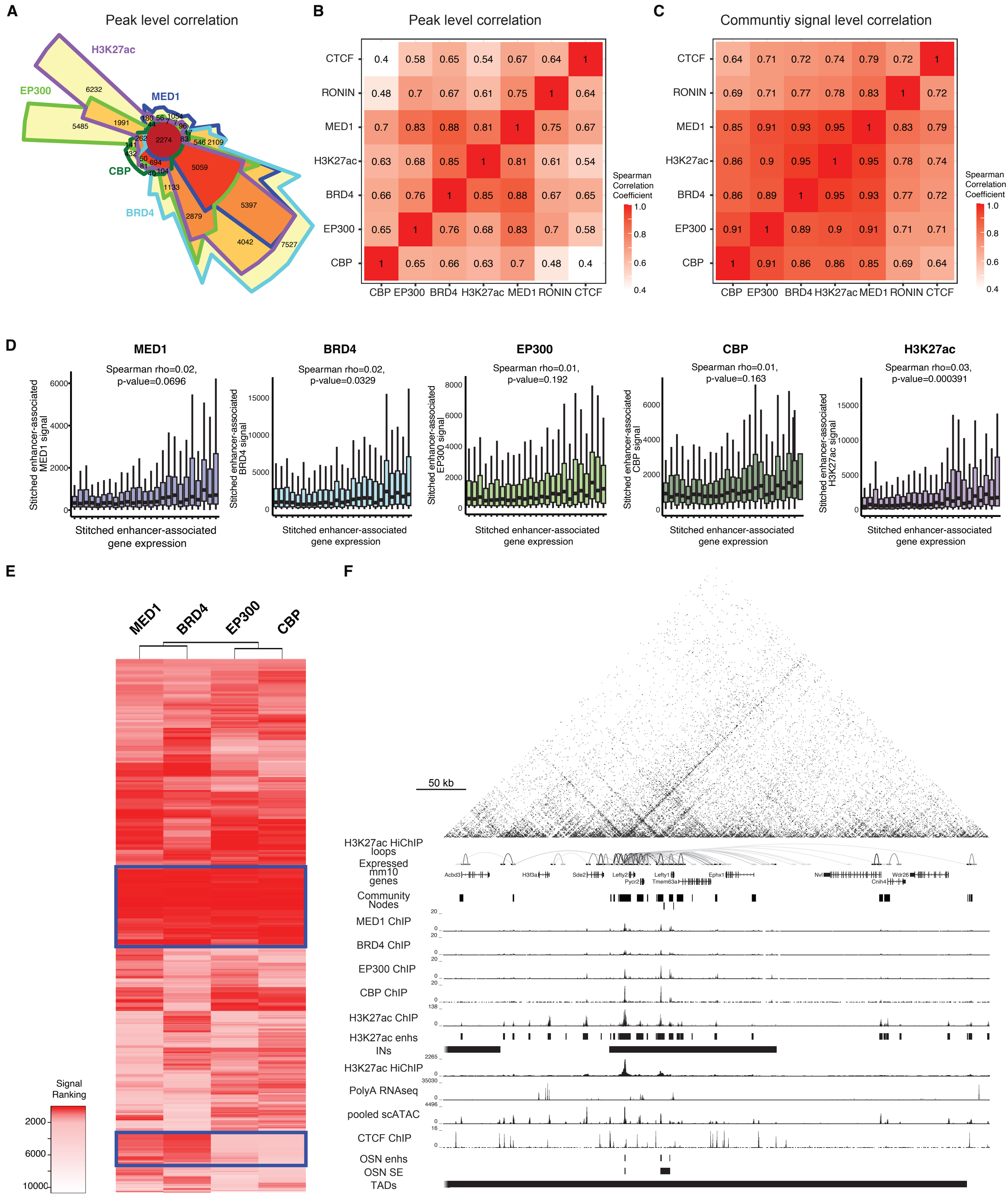

**Figure S3: The distributions of co-activator proteins are highly correlated.** (**A**) Chow-Ruskey diagrams showing overlap among ChIP-seq peaks for MED1, BRD4, EP300, CBP, and/or H3K27ac. Peak pairs were defined as overlapping when sharing at least one base pair (≥1 bp). The area of each section of the diagram is proportional to the number of peaks within it. (**B**) Heatmap showing pairwise Spearman correlation coefficients quantifying the similarity of protein distribution patterns across genome-wide peaks. (**C**) Heatmap showing pairwise Spearman correlation coefficients quantifying the similarity of protein distribution patterns across communities. (**D**) Distribution of ROSE-assigned per-stitched-enhancer-associated gene signal for MED1, BRD4, EP300, CBP, or H3K27ac, arranged into 25 groups by increasing gene expression level. The MED1 panel (left-most) is repeated from Fig. 1G and is provided here for ease of comparison. (**E**) Hierarchically clustered heatmap showing co-activator signal ranking at 3D-SEs. The top box highlights communities exhibiting uniformly high signal enrichment across all four co-activators. The bottom box shows communities with a differential enrichment pattern, characterized by high MED1 and BRD4 signals and relatively lower EP300 and CBP signals. (**F**) Genome browser view of the *Lefty1* community. The community nodes of the *Lefty1* community show greater levels of signal enrichment for some co-activators (EP300, CBP) than others (MED1, BRD4). Window size: 545 kb. ChIP-seq tracks are RPM normalized. The ends of TADs and insulated neighborhoods are faded to indicate that these features extend beyond the boundaries of this region.

Supplemental Figure 4

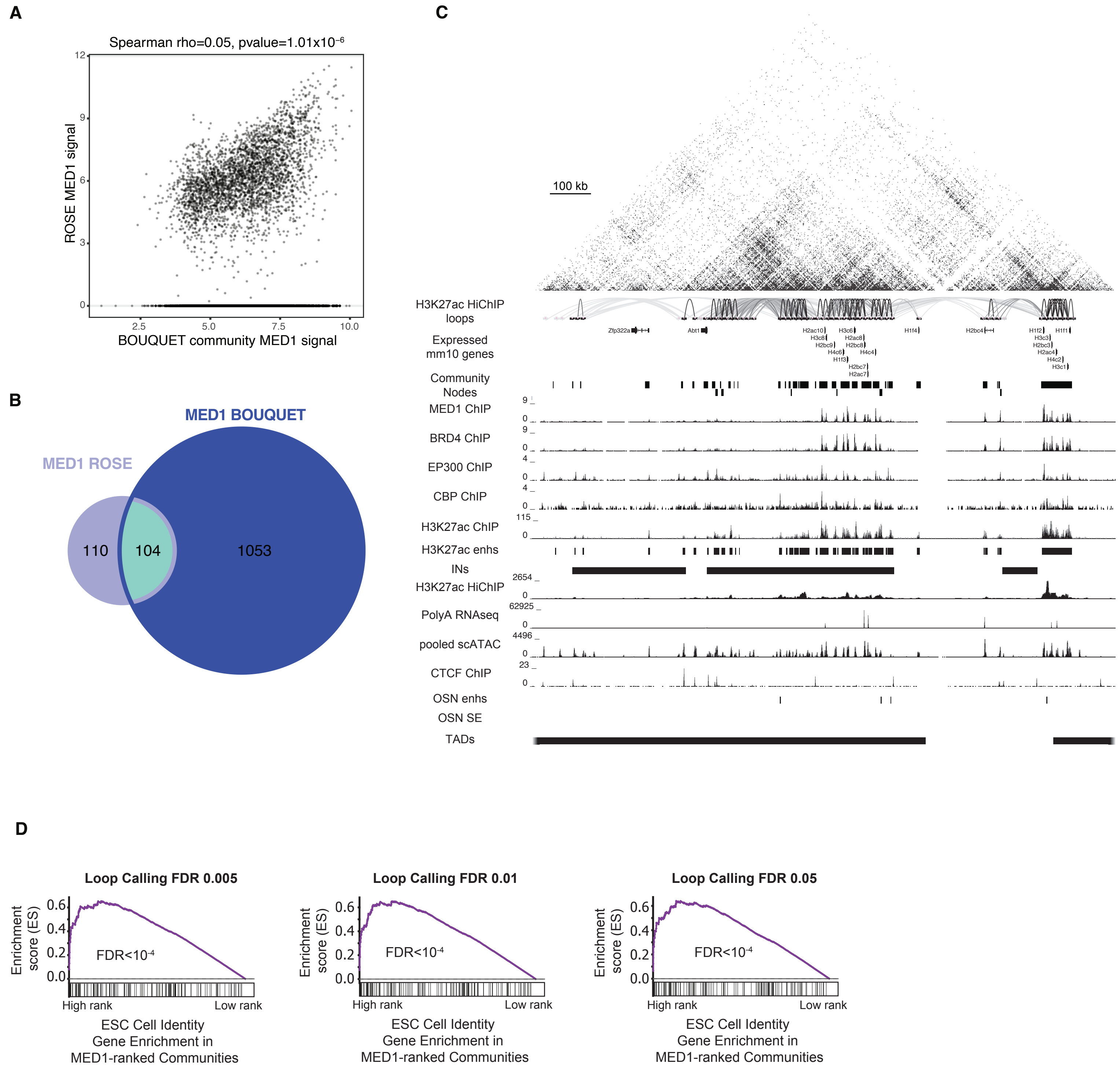

**Figure S4: Linear SEs and 3D-SEs capture partially distinct CRE sets per gene.** (**A**) Distribution of ROSE-assigned per-gene MED1 signal (y-axis) versus community-assigned per-gene MED1 signal (x-axis) (Spearman rho = 0.05). (**B**) Overlap between mESC linear SE-associated genes identified by the ROSE algorithm and 3D-SE–associated genes identified by BOUQUET. (**C**) Genome browser view of a histone gene cluster identified as a 3D-SE by BOUQUET in mESC. Histone genes are predominantly regulated by promoter elements; thus, their high levels of co-activator enrichment are overlooked by enhancer-based signal assignment methods such as ROSE (note the lack of super-enhancers in the OCT4/SOX2/NANOG super-enhancer ("OSN SE") track). ChIP-seq tracks are RPM normalized. (**D**) Pre-ranked gene set enrichment analysis (GSEA) for mESC cell identity genes across communities independently ranked by MED1 across three loop-calling thresholds. The mESC cell identify gene set derives from the "Embryonic Stem Cell" gene list in the PanglaoDB_Augmented_2021 database (see Supplemental Methods for additional details). Black tick marks denote individual mESC cell identity genes positioned by their community signal ranks.

Supplemental Figure 5

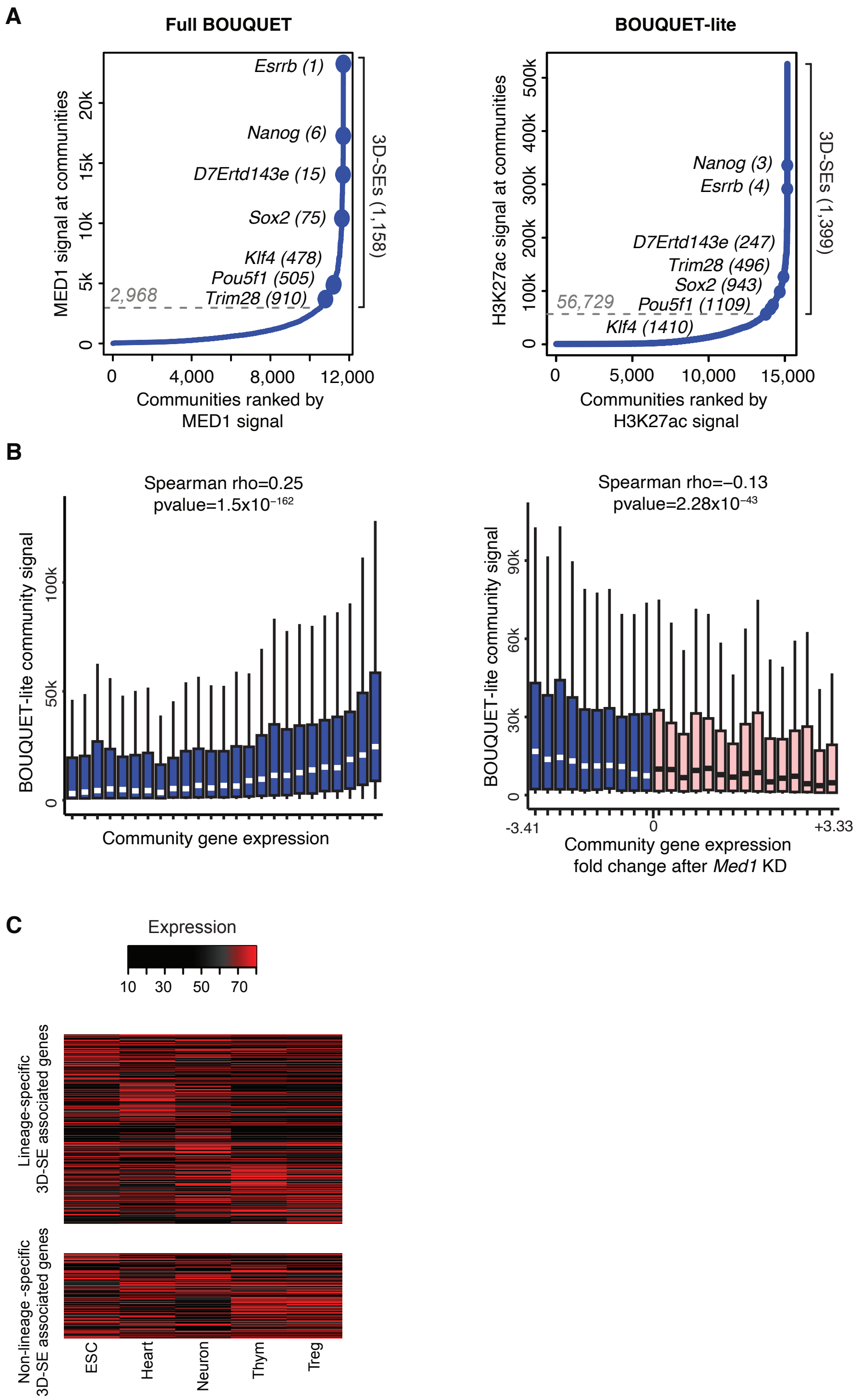

**Figure S5: BOUQUET-lite is a largely suitable alternative for full BOUQUET when data availability is limited.** (**A**) Left: Distribution of MED1 ChIP-seq signal across CRE communities identified by full BOUQUET. Right: Distribution of H3K27ac HiChIP signal across CRE communities identified by BOUQUET-lite. The total number of 3D-SEs is shown on the right of each graph. Dashed lines mark algorithmically determined signal cutoffs distinguishing 3D-SEs from moderate-signal communities. The MED1 ranking panel is reproduced from Fig. 2B for ease of comparison. (**B**) Left: Distributions of community-assigned per-gene H3K27ac HiChIP signal across grouped genes. Genes are sorted into 25 groups by increasing gene expression level. Right: Distributions of community-assigned per-gene H3K27ac HiChIP signal across grouped genes following knockdown of *Med1*. Genes are sorted into 25 groups by gene expression fold-change following *Med1* knock-down. (**C**) Heatmap showing expression of 3D-SE–associated genes across five mouse cell/tissue types. The top panel displays genes associated with 3D-SEs in only one cell/tissue type, and the bottom panel displays genes associated with 3D-SEs in multiple cell/tissue types. Expression values represent percent-scaled, rank-normalized TPM, where TPM values are ranked per sample and scaled to percentiles (0-100%), enabling relative expression visualization across samples independent of absolute levels.

Supplemental Figure 6

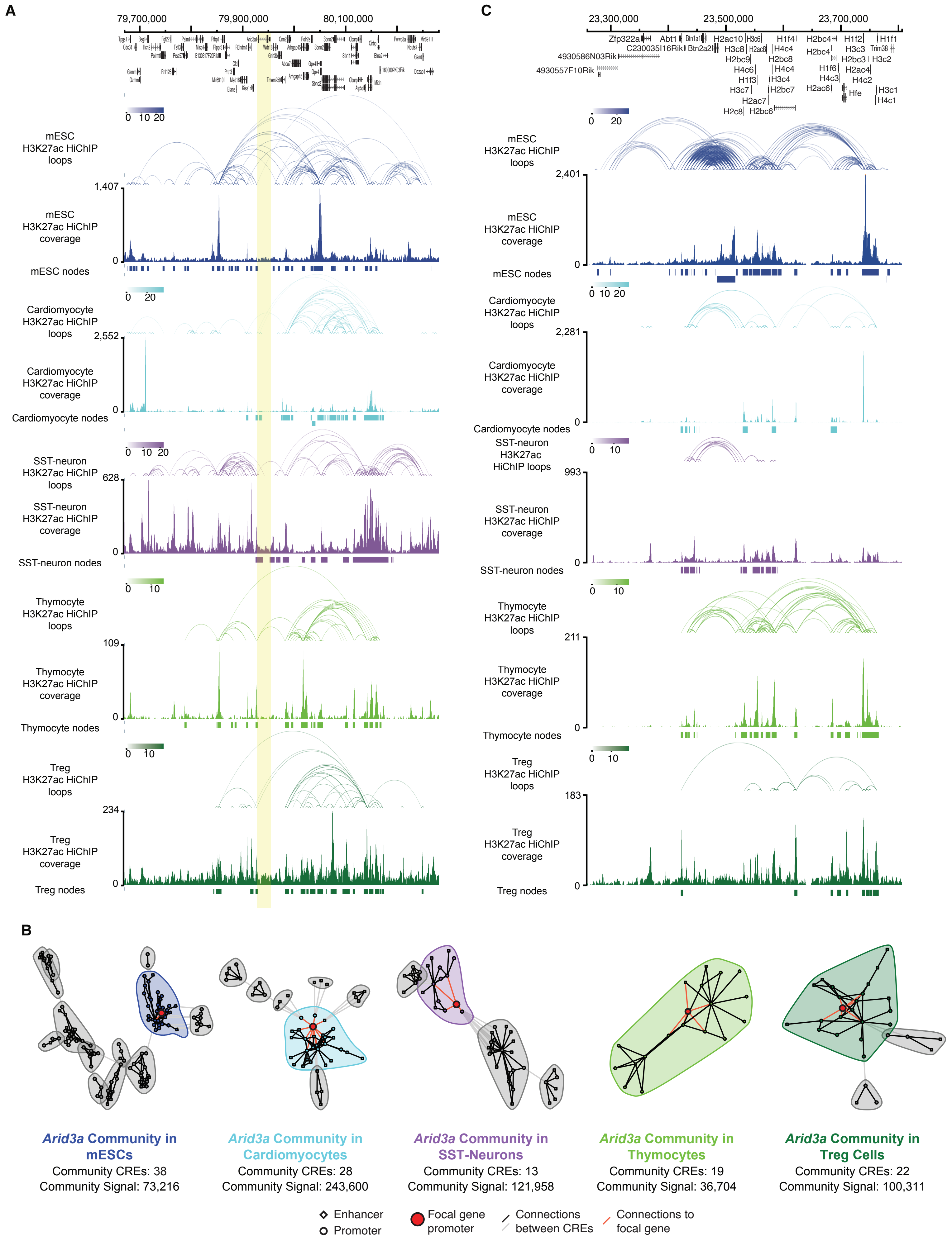

**Figure S6: 3D-SEs are largely cell-type-specific.** (**A**) ProteinPaint genome browser visualization of the *Arid3a* CRE community, identified as a gene associated with a 3D-SE in all five mouse cell/tissue types shown. The location of *Arid3a* is highlighted in yellow. Tracks show high-confidence H3K27ac HiChIP loops and H3K27ac HiChIP coverage. The CREs (i.e. nodes) contained in the *Arid3a* 3D-SE community are shown for each cell type/tissue. Window size: 610 kb (chr10: 79,670,462 - 80,281,141, mm10). (**B**) Network diagrams of the *Arid3a* components in each of five cell/tissue types. The *Arid3a* community is marked by the colored cloud in each cell type; gray clouds mark other communities within the component that do not contain *Arid3a*. The *Arid3a* focal gene promoter is marked by a red circle. (**C**) ProteinPaint genome browser visualization of histone gene cluster communities identified as 3D-SEs in the five mouse cell/tissue types shown. Tracks show high-confidence H3K27ac HiChIP loops and H3K27ac HiChIP coverage. The CREs nodes contained in each histone gene cluster 3D-SE community are shown for each cell type/tissue. Window size: 547 kb (chr13: 23,259,392 - 23,806,396, mm10).

Supplemental Figure 7

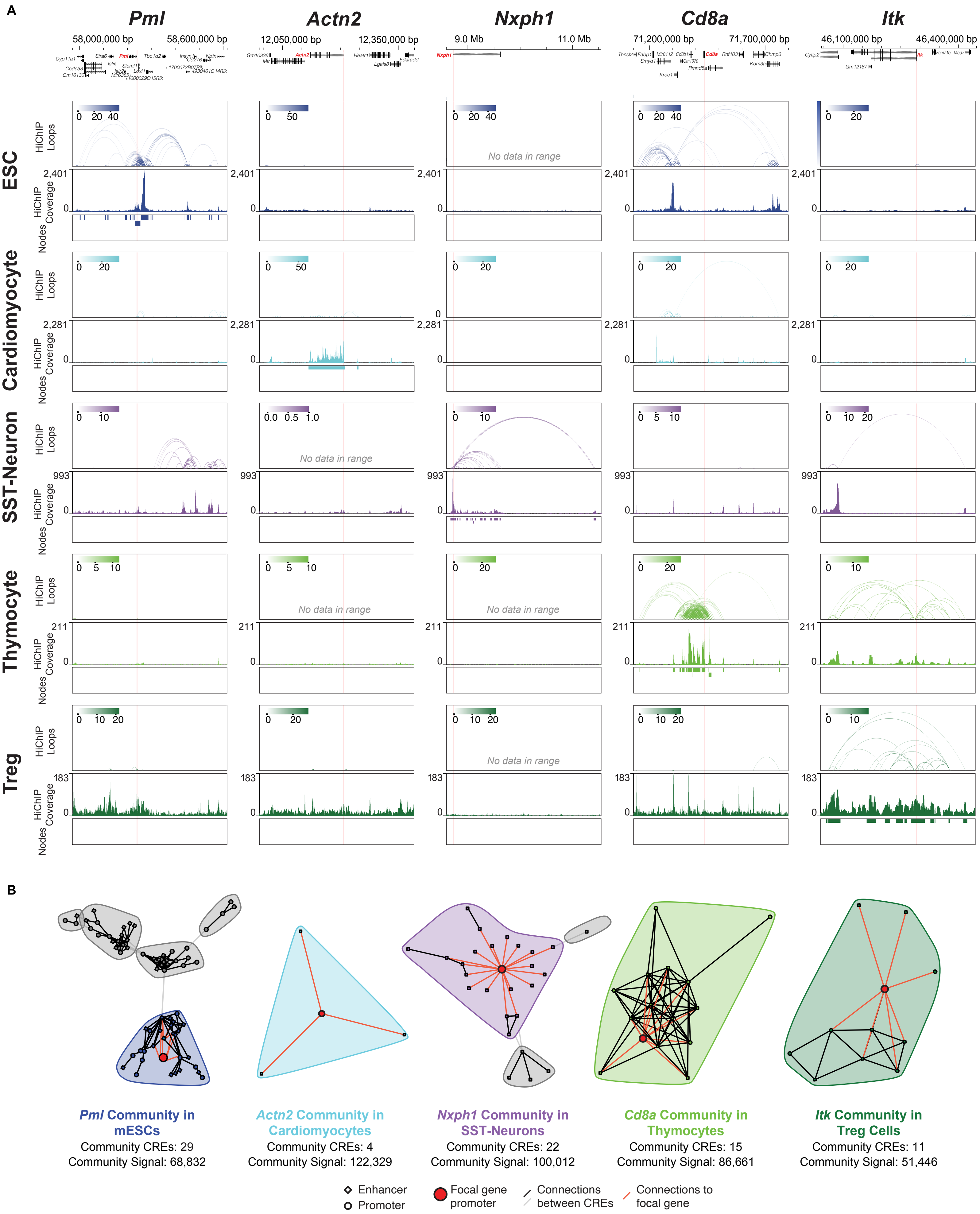

**Figure S7: Communities of cell-identity-defining genes are cell-type-specific.** (**A**) ProteinPaint genome browser tracks of 3D-SE–associated cell-identity-defining genes in five mouse cell/tissue types. Tracks show high-confidence H3K27ac HiChIP loops and H3K27ac HiChIP coverage (mm10 coordinates: *Pml*: chr9: 57,982,351 - 58,618,168; *Actn2:* chr13: 12,160,949 - 12,489,890; *Nxph1*: chr6: 8,910,656 - 9,881,733; *Cd8a*: chr6: 71,134,802 - 71,663,053; *Itk*: chr11: 46,295,139 - 46,446,895). The CRE nodes within each 3D-SE community are shown for their defining cell type/tissue. The name of each cell-identity-defining gene is marked in red in the gene track (*top*), and its TSS is marked by a vertical red line spanning all browser tracks. (**B**) Network diagrams of the component of each example cell-identity-defining gene in each of five cell/tissue types (ref: panel A). The community of each cell-identity-defining gene is marked by the colored cloud in its defining cell type/tissue; gray clouds mark other communities within the component that do not contain the cell-identity-defining gene of interest. The focal gene promoter is marked in each diagram by a red circle.

Supplemental Figure 8

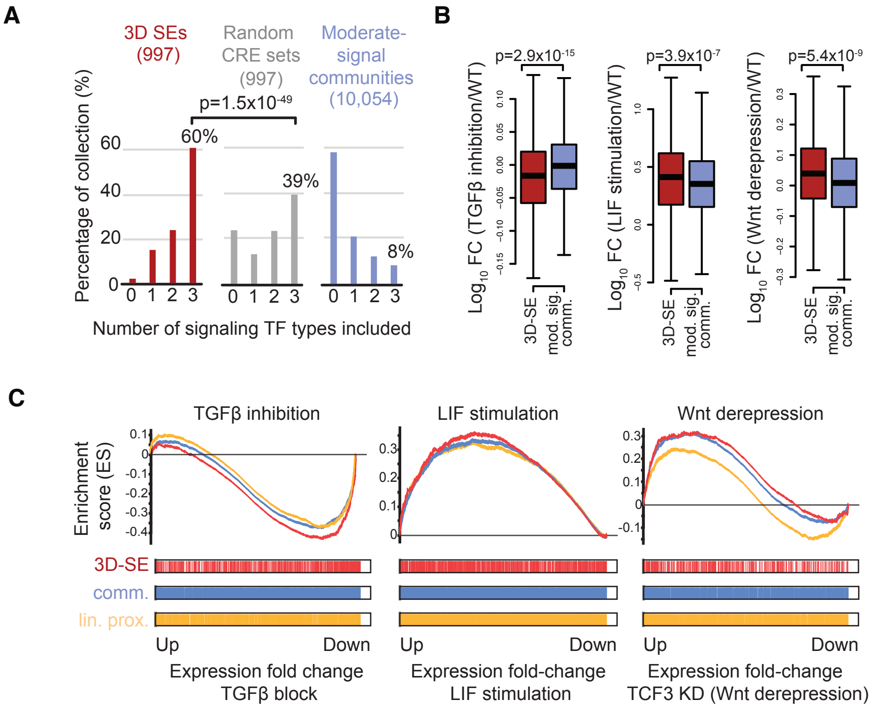

**Figure S8: CRE communities accurately predict signaling pathway perturbation.** (**A**) Distribution of binding sites for three signaling transcription factors (TFs) (STAT3, SMAD3, and TCF3) at CREs within different categories of community. Bars denote the percentage of communities in each category with the presence of binding sites for 0, 1, 2, or all 3 types of TFs. Here the binding sites within 3D-SEs (red, 997 communities), randomized sets of CREs with the same CRE count distribution as the 3D-SEs (grey, 997 randomized sets), and moderate-signal CRE communities (blue, 10,054 communities) are compared. (**B**) Change in expression of 3D-SE genes (red) and moderate-signal community genes (blue) following perturbation of the TGF-β pathway (*left*), stimulation of the LIF pathway (*middle*), and perturbation of the Wnt pathway (*right)*. N.B., the Wnt pathway largely represses gene expression in wild-type mESC, and, correspondingly, the trend of gene expression following Wnt pathway perturbation is up-regulation. (**C**) Gene set enrichment analysis (GSEA) of gene expression changes after manipulation of the TGF-β, LIF, and Wnt pathways. "3D-SE" (red), "community" (blue), and "linear prox" (gold) indicate genes in 3D-SE communities with a signaling-TF-bound CRE, genes in all communities with a signaling-TF-bound CRE, and genes that were the most linearly proximal to a signaling-TF-bound CRE, respectively.

Supplemental Figure 9

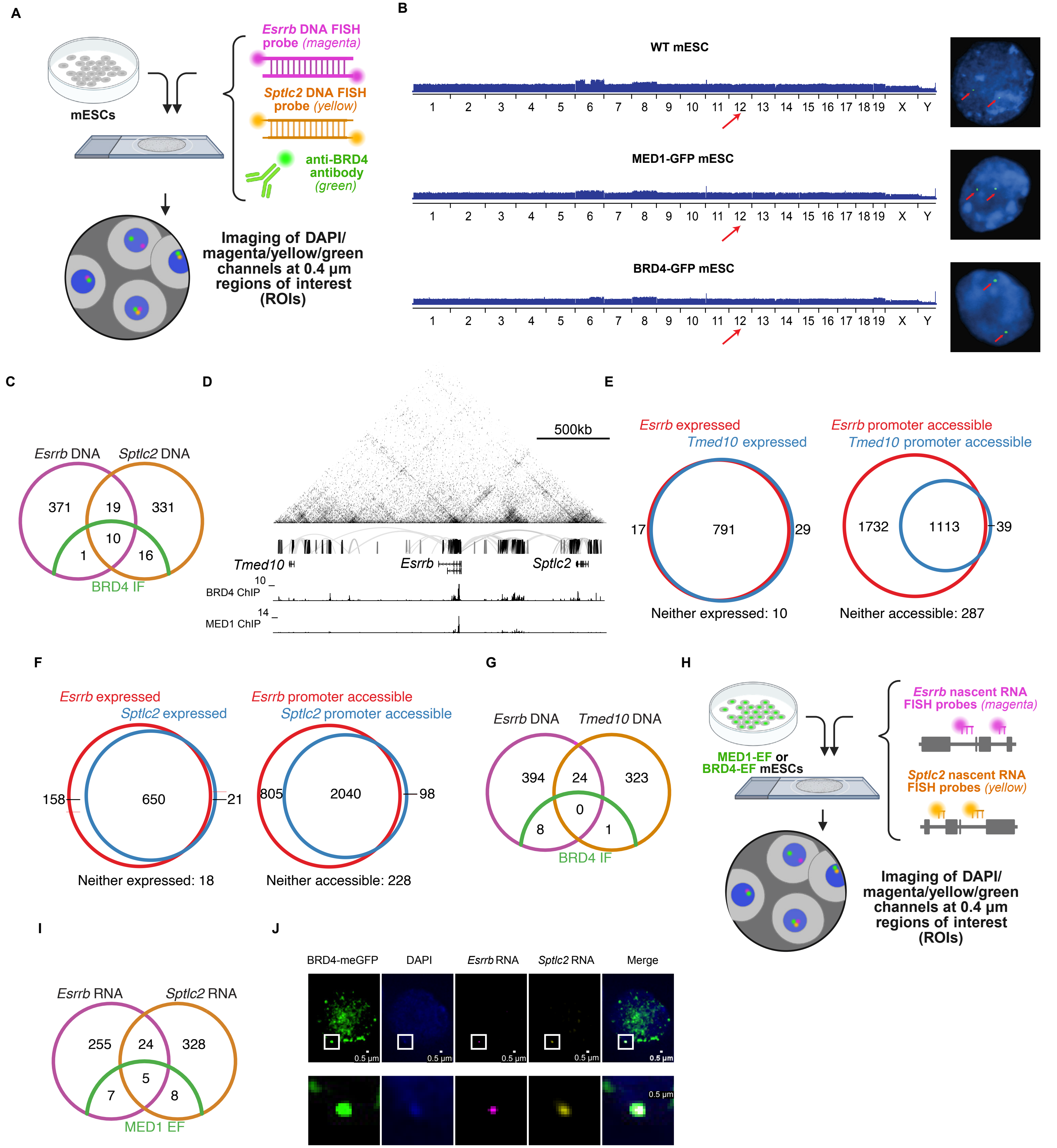

**Figure S9: Fluorescence microscopy of co-localized 3D-SE components.** (**A**) Experimental workflow of DNA fluorescence *in situ* hybridization (FISH) with BRD4 immunofluorescence (IF) in mESCs. Created in BioRender. Maher, K. (2025) https://BioRender.com/jkj4wow. See Supplemental Methods for additional details. (**B**) Left panels: IGV tracks showing whole-genome sequencing read coverage across mouse chromosomes in wild-type mESCs (top), endogenously MED1-tagged mESCs (middle), and endogenously BRD4-tagged mESCs (bottom). Right panels: Fluorescence microscopy of mESC nuclei with probes targeting chromosome 12 (host of DNA/RNA FISH target community containing *Esrrb*/*Sptlc2* and *Tmed10*) in wild-type mESCs (top), endogenously MED1-tagged mESCs (middle), and endogenously BRD4-tagged mESCs (bottom). Blue = DAPI stain for chromatin; green = DNA probes for chromosome12. Images represent a single z-plane. (**C**) Venn diagram showing the overlap of regions of interest (ROIs) containing signal for *Esrrb* DNA, *Sptlc2* DNA, and BRD4 IF puncta in various combinations. (**D**) UCSC Genome Browser view of the locus containing *Tmed10/Esrrb/Sptlc2*, the targets of the DNA FISH experiments. Window size: 2.3Mb (chr12: 85,156,077-87,439,412, mm10). ChIP-seq tracks are RPM normalized. (**E**) Left panel: Venn diagram showing the expression of *Esrrb* and *Tmed10* in single mESCs via scRNA-seq. Individual mESCs with >0 mapped reads for a given gene are defined as expressing that gene. Right panel: Venn diagram showing the accessibility of the *Esrrb* and *Tmed10* promoters in single mESCs, via scATAC-seq. Individual mESCs with >0 mapped reads within the 4 kb region surrounding each gene's TSS are defined as having an accessible promoter for that gene. (**F**) Left panel: Venn diagram showing the expression of *Esrrb* and *Sptlc2* in single mESCs via scRNA-seq. Right panel: Venn diagram showing the accessibility of the *Esrrb* and *Sptlc2* promoters in single mESCs, via scATAC-seq. (**G**) Venn diagram showing the overlap of regions of interest (ROIs) containing signal for *Esrrb* DNA, *Tmed10* DNA, and BRD4 IF puncta in various combinations. (**H**) Experimental workflow of RNA FISH in mESCs with either endogenously tagged MED1 or endogenously tagged BRD4. Created in BioRender. Maher, K. (2025) https://BioRender.com/b9h2fpm. See Supplemental Methods for additional details. (**I**) Venn diagram showing the overlap of ROIs containing signal for nascent *Esrrb* RNA, nascent *Sptlc2* RNA, and MED1 EF puncta in various combinations. (**J**) Fluorescence microscopy of mESC nuclei following RNA FISH with BRD4 EF. Blue = DAPI stain for chromatin; magenta = nascent RNA probes for *Esrrb*; yellow = nascent RNA probes for *Sptlc2*; green = BRD4 EF. Images represent a single z-plane. Scale bar for top panels = 0.5 µm. Lower panels are zoomed-in views of the same ROI shown in the top panels.

**Supplementary Data: Table of Contents**

**Data S1:** All CRE nodes defined in full BOUQUET for mESCs. Each CRE can be a promoter (Plable=1) or an enhancer (Elabel=1 and Plabel=0). For an enhancer CRE, both gene_symbol and RefSeq ID show "enhancer." Related to Supplemental Figure 1.

**Data S2:** All CRE pairs (edges) supported by HiChIP loops and/or INs for mESCs. For type column (3rd), 0=supported by Loop only; 1=supported by IN only; 2: supported by both Loop and IN. Related to Supplemental Figure 1.

**Data S3:** H3K27ac HiChIP loops called by Hi-LOW (using FitHiChIP, FDR<0.01), with putative anchors defined as the merged set of H3K27ac ChIP-seq peaks and promoters (+/- 2 kb around the TSS). These loops are used as input for full BOUQUET for mESCs. The $7^{th}$ column is number of HiChIP read pairs supporting each loop. Compare to Data S7. Related to Supplemental Figure 1.

**Data S4:** Insulated Neighborhoods called in mm10 for mESCs, used as input for full BOUQUET. Related to Supplemental Figure 1.

**Data S5:** Full BOUQUET output table for mESCs. Each entry represents a community built off of a single focal gene. The Community.Enhancer column reports counts for CREs overlapping either H3K27ac ChIP-seq peaks or promoters (left) versus CREs overlapping only H3K27ac ChIP-seq peaks (right). Related to Figure 1, 2, Supplemental Figure 1, 2, 4.

**Data S6:** Full BOUQUET 3D-SE gene lists ranked by ChIP-seq signal for specified factors. For each factor, two versions of the 3D-SE gene list are provided: the original condensed 3D-SE gene list with unique promoters (left) and the expanded 3D-SE gene list (right) that includes all 3D-SE gene isoforms. Related to Figure 2.

**Data S7:** H3K27ac HiChIP loops called by Hi-LOW (using FitHiChIP, FDR<0.01), with putative anchors defined as the merged set of H3K27ac HiChiP peaks and promoters (+/- 2 kb around the TSS). These loops are used as input for BOUQUET-lite in mESCs. Compare to Data S3. Related to Figure 3, Supplemental Figure 5.

**Data S8:** All CRE nodes defined in BOUQUET-lite for mESCs. Each CRE can be a promoter (Plable=1) or an enhancer (Elabel=1 and Plabel=0). For enhancer CRE, both gene_symbol and RefSeq ID shows "enhancer." Related to Figure 3, Supplemental Figure 5.

**Data S9:** All CRE pairs (edges) supported by HiChIP loops by BOUQUET-lite for mESCs. For type column (3rd), 0=supported by Loop only. Related to Figure 3, Supplemental Figure 5.

**Data S10:** BOUQUET-lite output table for mESCs. Each entry represents a community built off of a single focal gene. The Community.Enhancer column reports counts for CREs overlapping either H3K27ac HiChIP peaks or promoters (left) versus CREs overlapping only H3K27ac HiChIP peaks (right). Related to Figure 3, Supplemental Figure 5.

**Data S11:** BOUQUET-lite output table for cardiomyocytes. Each entry represents a community built off of a single focal gene. The Community.Enhancer column reports counts for CREs overlapping either H3K27ac HiChIP peaks or promoters (left) versus CREs overlapping only H3K27ac HiChIP peaks (right). Related to Figure 3.

**Data S12:** BOUQUET-lite output table for SST-neurons. Each entry represents a community built off of a single focal gene. The Community.Enhancer column reports counts for CREs overlapping either H3K27ac HiChIP peaks or promoters (left) versus CREs overlapping only H3K27ac HiChIP peaks (right). Related to Figure 3.

**Data S13:** BOUQUET-lite output table for thymocytes. Each entry represents a community built off of a single focal gene. The Community.Enhancer column reports counts for CREs overlapping either H3K27ac HiChIP peaks or promoters (left) versus CREs overlapping only H3K27ac HiChIP peaks (right). Related to Figure 3.

**Data S14:** BOUQUET-lite output table for regulatory T (Treg) cells. Each entry represents a community built off of a single focal gene. The Community.Enhancer column reports counts for CREs overlapping either H3K27ac HiChIP peaks or promoters (left) versus CREs overlapping only H3K27ac HiChIP peaks (right). Related to Figure 3.

**Data S15:** BOUQUET-lite 3D-SE gene lists. For each cell type, two versions of the 3D-SE gene list are provided: the original condensed 3D-SE gene list with unique promoters (left) and the expanded 3D-SE gene list (right) that includes all 3D-SE gene isoforms. Related to Figure 3, Supplemental Figure 5.

**Data S16:** DESEQ2 outputs for differential expression analysis between WT and D7Ertd143e enhancer-deletion RNA-seq in mESCs. Related to Supplemental Figure 2.

**Data S17:** TPM values from WT and D7Ertd143e enhancer-deletion RNA-seq in mESCs. Related to Supplemental Figure 2.

**Data S18:** Linear stitched enhancers by MED1 signal with super-enhancer/typical enhancer annotation defined by the ROSE algorithm. Related to Figure 1, Supplemental Figure 3, 4.

**Data S19:** TPM values from WT and Med1 knockdown RNA-Seq and expression fold changes upon Med1 knockdown in mESCs. Related to Figure 1, Supplemental Figure 2, 5.

**Data S20:** Linear stitched enhancer regions identified using the ROSE algorithm, annotated as super-enhancers or typical enhancers based on BRD4, CBP, EP300, and H3K27ac ChIP-seq signal. Related to Supplemental Figure 3.

**Data S21:** Coefficient of variance (CoV) among MED1, BRD4, EP300, and CBP signal across full BOUQUET communities. Related to Supplemental Figure 3.

**Data S22:** Top 10 ranked PanglaoDB_Augmented_2021 terms for Gene Ontology (GO) analyses. Related to Figures 2, 3.

**Data S23:** Gene sets used for Gene Set Enrichment Analyses (GSEAs). Related to Figures 2, 3, Supplemental Figures 4, 5, 8.

**Data S24:** TPM values from WT and LIF stimulated mESC RNA-seq and expression fold changes upon LIF stimulation (generated by re-analyzing polyA-selected RNA-seq of WT and LIF stimulated mESC from E-MTAB-1796). Related to Supplemental Figure 8.

**Data S25:** Public datasets used in this study.

**Data S26:** Nascent RNA probes for RNA FISH experiments. Related to Figure 4, Supplemental Figure 9.

#### Supplemental Materials and Methods

##### Computational analysis of ChIP-seq data

Published ChIP-seq datasets from NCBI GEO for MED1 (GSM560348), BRD4 (GSM3084073), EP300 (GSM918750), CBP (GSM1246866), H3K27ac (GSM1526287), RONIN (GSM1246868), H3K4me3 (GSM1526288), CTCF (GSM747534), H3K27me3 (GSM1526291), OCT4 (GSM1082340), SOX2 (GSM1082341), NANOG (GSM1082342), TCF3 (GSM307143, GSM307142), STAT3 (GSM686673), and SMAD3 (GSM539542, GSM539541) in mouse Embryonic Stem Cells (mESCs) were processed to find statistically enriched regions, i.e. peaks, and quantify coverage (Supp. Data S25). Quality control, mapping, coverage track-building, and peak calling were performed with the SEASEQ pipeline (v3.0) (1). Briefly, reads were mapped to the mouse reference genome sequence version mm10 (GRCm38). To identify narrow enriched regions (peaks) for H3K27ac, MED1, BRD4, EP300, CBP, RONIN, CTCF, H3K4me3, TCF3, SMAD3, and STAT3, MACS (v1.4.2) (2) was utilized through SEASEQ with the following parameters: keep-dup= auto, band width = 300, model fold = 10,30, p-value cutoff = 1.00e-09. If corresponding control libraries existed, they were used in peak-calling and are noted in Supplemental Data S25. To identify significantly enriched regions of H3K27me3, we used broad peak finding within SEASEQ. Briefly, broad peak finding in SEASEQ utilizes SICER (v1.1) (3) with the following parameters: W=200; G=400 and FDR=0.01.

##### Computational analysis of RNA-seq data

For our reference gene annotation, we downloaded RefSeq Release 108 for GRCm38.p6 from the NCBI FTP site, https://ftp.ncbi.nlm.nih.gov/refseq/M_musculus/annotation_releases/108/GCF_000001635.26_GRCm38.p6/GCF_000001635.26_GRCm38.p6_genomic.gtf.gz. In order to focus on high-confidence gene annotations, we filtered 1) *against* genes annotated as “unknown transcript,” 2) *against* MGI-predicted genes, i.e. those whose gene names start “Gm,” and 3) *for* genes whose identifiers start with “NM” or “NR”, which are curated protein-coding and non-protein-coding genes, respectively, resulting in 40,618 genes for downstream analysis.

RNA-seq reads were mapped to the mm10 reference genome to which the sequences of ERCC spike-in probes were added using hisat2 (v2.1.0) (4) with default parameters. Htseq-count (v0.11.2) (5) was used with parameters “--stranded=reverse -m intersection-strict –gene_id” to count reads mapped to any transcript of each gene. For each gene, we extracted its total non-redundant, i.e. collapsed, exon length using the merge function of bedtools v2.30.0 (6) and used this collective exon length to calculate transcript per million (TPM) values. TPM log2 fold-changes were calculated after adding a pseudocount of 1 to each TPM. For differential expression analysis between samples of different experimental conditions, DESeq2 (7) was used to calculate adjusted p-values for expression changes of individual genes using read counts.

In wild type V6.5 mESCs (GSM1246869) we determined the expression level of each gene using transcript per million (TPM) values by analyzing polyA-selected RNA-seq as above (8).

#### Identification of linear super-enhancers

ROSE (https://bitbucket.org/young_computation/rose/) was used to identify linear super-enhancers in mESCs by generally following a protocol from (9). In the previous work, the mm9 reference genome was used, and we instead used mm10, so we formatted our reference GTF from above for ROSE input. We acquired OCT4, SOX2, NANOG, MED1, BRD4, EP300, CBP, H3K27ac ChIP-seq and corresponding inputs. To more closely represent the analysis performed in previous SE studies, we realigned the raw ChIP-seq reads to the mm10 reference genome using bowtie (10) with parameters -k 1 -m 1 -n 2 --best and -l set to the read length. Peaks of OCT4, SOX2, and NANOG were identified using MACS (v1.4.2) with corresponding control, -g mm, -p 1e-9. The intersection of these peaks was determined using bedtools intersect and used as input for ROSE alongside using MED1, BRD4, EP300, CBP, or H3K27ac signal to rank stitched enhancers, corresponding input control, and with parameter -s 12500. The CLOSEST_GENE identified, i.e. the single gene whose transcription start site was most proximal to the center of a given stitched enhancer, was assigned to that stitched enhancer. N.B. that "signal" in the context of ROSE comes from reads-per-million-normalized background-subtracted density values that are expanded into signal based on the width of the stitched enhancer in question. In downstream analysis where ROSE signal and BOUQUET signal are compared directly, how those signals are calculated are not fully in alignment.

#### HiChIP/HiC Looping Organization Workflow (Hi-LOW): one-click identification of high-confidence HiChIP loops

##### Step 1: Initial processing of HiChIP data

H3K27ac HiChIP data from wild-type V6.5 mouse Embryonic Stem Cells (mESCs) were downloaded from the Gene Expression Omnibus (GEO) database (GSE99519, Supp. Data S25) (11). Reads from three replicates were pooled, though each sample gave similar results when processed individually. We developed Hi-LOW (github.com/stjude/HiLOW) to facilitate the one-click basic analysis of HiChIP data from FASTQ to loops by wrapping existing tools. Hi-LOW executes HiC-Pro (v 2.11.1) (12) to perform read alignment, read filtering, quality check, and contact matrix building using default parameters, unless otherwise specified below. To achieve improved performance, Hi-LOW splits the input FASTQ files into multiple files, each with 10 million read-pairs, which were processed by HiC-Pro in parallel mode. HiC-Pro was configured so that each end of the read-pairs was mapped independently to the mm10 reference genome using bowtie2 (v2.3.5.1) (13). For global and local alignment, we used the options "bowtie2 --very-sensitive -L 30 --score-min L,-0.6,-0.2 --end-to-end –reorder" and "bowtie2 --very-sensitive -L 20 --score-min L,-0.6,-0.2 --end-to-end –reorder" respectively. For read mapping, we used MAPQ threshold=10 and removed both multi-mapped reads (RM_MULTI=1) and duplicated read-pairs (RM_DUP=1). Taking the final valid read-pairs generated from HiC-Pro as input, we used the "hicpro2juicebox.sh" utility script to create a HiChIP contact map in ".hic" format, which was used for downstream analysis.

**Step 2: Loop calling using HiChIP data**

Predefined loop anchors are required for downstream loop-calling steps. In mESCs, we used the collapsed union of active enhancers and promoters, defined by H3K27ac ChIP-seq peaks and promoters, both defined as above. The collapsed union of these regions was generated using the merge function of bedtools v2.30.0 was used as a set of candidate loop anchors.

Hi-LOW uses FitHiChIP (v9.0) (14) to determine statistically significant pairwise interactions/loops in HiChIP datasets. The valid HiChIP read-pairs generated by HiC-Pro and our candidate loop anchors were used as input for FitHiChIP. Briefly, the genome was divided into 5 kb bins, for which read coverage was quantified. For the identification of loops, 5 kb bins that overlapped candidate loop anchors were determined by FitHiChIP. In this study, we required each significant loop to connect bins where at least one end overlapped a candidate loop anchor (Peak-To-All mode). In addition, we used a loose background estimation (FitHiChIP(L) mode) to infer a null model, i.e. expected contact distribution, from HiChIP read-pairs. The binomial distribution was used in FitHiChIP on the observed contact frequencies to estimate the *P*- values, which were then corrected for multiple testing. Interactions having false discovery rate (FDR) < 0.01 were considered significant and were reported as loop calls. Subsequently, the loops were formatted as BEDPE for downstream visualization and CRE association. To examine the impact of loop-calling FDR thresholds on BOUQUET results, loop calling was repeated with two additional thresholds, one more stringent (FDR 0.005) and one more permissive (FDR 0.05), relative to the default threshold (FDR 0.01). For downstream community detection and unless otherwise stated, loops called at FDR<0.01 were used.

#### Insulated neighborhood (IN) calling

**Step 1: ChIA-PET data processing**

Raw mESC SMC1A ChIA-PET datasets (GSE57911) were processed using an in-house bioinformatics pipeline derived as strictly as possible from a previously reported strategy to enable close comparison with previous results and now shared at https://github.com/stjude/Dowen_Fan (15). Each read-pair from each replicate of ChIA-PET data was examined for the occurrence(s) of one or both linker sequences (Linker A=CTGCTGTCCG; Linker B=CTGCTGTCAT) at any position, and the distribution of the linker positions in reads was examined. Consistent with the prior report and the activity of ECOP151, most of the linkers in end-reads that contained them were located at around 27 bp from the 5′-end. For end-reads with a single perfect linker match, i.e. no mismatches and no indels, each end-read was trimmed at the 5′ position of this perfect linker sequence hit (cutadapt v4.3 (16) with parameters --no-indels --pair-adapters --pair-filter any -m 27:27 -M 36:36 -n 1 -O 10 -e 0). Read-pairs comprising trimmed end-reads between 27 bp-36 bp were separated into the following categories: chimeric, i.e. read-pairs comprised of end-reads where one end-read has one A linker and the other end-read has one B linker; non-chimeric, i.e. read-pairs comprising end-reads where both contain the same A or B linker; or other, i.e. read-pairs with other linker configurations. Non-chimeric reads are preferred, as they are thought to represent interactions that occur in the cells rather than during the experimental preparation.

Using Bowtie (v 1.2.2), non-chimeric read-pairs were mapped onto the mouse genome reference version mm10 in single-end mode with parameters –k 1, -v 1, -m 1 –e 70 –best –strata. For read-pairs where each end-read successfully aligned, samtools (v 1.15.1) merge (-@ 16), sort (-@ 16), and fixmate were used to re-mate and check proper pairing.

For peak calling, the remaining, aligned read-pairs were separated into a set of end-reads and further processed. For connectivity analysis, presumed PCR duplicate read-pairs, i.e. with identical chromosome, start, and end coordinates for each end-read, were identified, de-duplicated, and further processed as described below.

Surviving reads from both replicates were merged for peak calling. Peaks were called using MACS (v1.4.1) (2) with parameters -p 1e-9, --keep-dup=2, nomodel and nolambda, and these peaks were considered candidate loop anchors. PETs thought to represent self-ligation events were removed if their end-reads were separated by less than 5 kb. Connectivity between pairs of candidate loop anchors was quantified by overlapping PETs with the anchor peaks, defined earlier with MACS, using bedtools (v 2.30.0) (pairToBed -f 0.75). As used previously (15), significant interactions between two candidate loop anchors were required to meet two simultaneous cutoffs: that they were connected by >=3 PETs in the combined replicate data and that the FDR <=0.01 from a hypergeometric test (dhyper module in R, comparing the number of PETs connecting this anchor-pair against the total number of PETs connecting any anchor-pair and the number of PETs overlapping each anchor).

**Step 2: Insulated Neighborhood (IN) calling**

Insulated neighborhoods (INs) were defined as the significantly interacting SMC1A ChIA-PET loops (described above) whose anchors overlapped with CTCF ChIP-seq peaks called using SEASEQ and contained at least one gene. To identify the subset of SMC1A ChIA-PET loops that represent CTCF-CTCF interactions, we selected loops whose anchor regions both overlapped with CTCF ChIP-seq peaks by at least 1 bp using PairToBed (6). CTCF-CTCF interactions that contained at least one gene (Refseq 108, filtered as defined above) were derived using PairToBed -ospan in bedtools, selecting loops that spanned at least one entire gene, i.e. start1 of anchor1 < start of gene and end2 of anchor2 > end of gene. After removing the top 3 largest INs (> 6Mb) that are substantially larger than the rest of INs (< 3Mb), 20,240 INs were kept for downstream analysis.

#### Signaling transcription factor overlap with CRE communities

To explore the relationships between CRE communities and signaling transcription factors, each CRE community was first decomposed into a list of its constituent CREs. The binary presence/absence of an overlap (>1 bp) for TCF3, SMAD3, and/or STAT3 transcription factor ChIP-seq peaks was determined across all constituent CREs of a community, and then aggregated to determine whether each community contained a CRE that was bound by 0-3 types of signaling TF (Supp. Fig. 8A). We compared percentages of 0-bound, 1-bound, 2-bound, and 3-bound communities across three categories: 3D-SEs, moderate-signal communities, and randomized CRE sets as a control. Considering that 3D-SEs generally contained larger numbers of individual CREs, CRE sets made of the same number of CREs as 3D-SEs but comprising randomly selected CREs from moderate-signal communities were created to control for the potential enrichment effect of larger communities and allow for a fairer comparison between the two community types. Specifically, random CRE sets were created following the same distribution of CRE number per 3D-SE community. Using the binary TF overlap list generated above for all CREs, the 0-3 count of bound TFs was created for each random CRE set. This process was repeated 10,000 times to determine the percentage of simulated CRE-sets bound by 0, 1, 2, or 3 types of TFs (Supp. Fig. 8A, in gray). We observed a stable distribution of randomized community sets that fell into these categories (not shown). The chi-square significance calculation between real and simulated communities compared their distribution of community numbers across 4 categories: communities bound by zero, one, two, or all three types of TFs in a 2x4 table.

#### Genome-wide expression changes after perturbation (bulk RNA-seq)

Raw RNA-seq data were acquired from wild-type and *Med1* knock-down mESCs (GSM5399220) (17). TPMs were calculated for each of the two replicates separately (see method for RNA-Seq analysis). Fold changes of *Med1* KD/WT conditions were calculated after averaging TPMs across replicates for each condition and transforming by log2 after adding a pseudocount of 1 (Supp. Data S19).

Raw RNA-Seq data were acquired from mESCs treated with LIF for 0h (control) and 1h (LIF stimulation) (E-MTAB-1796 Arrayexpress dataset (18)). TPMs were calculated for each of the two replicates of (see method for RNA-Seq analysis). Fold changes of LIF stimulation/control conditions were calculated after averaging TPMs across replicates for each condition and transforming by log2 after adding a pseudocount of 1 (Supp. Data S24).

Precomputed gene expression changes after blocking TGF-β signaling by the inhibitor SB431542 were downloaded from a previous study (Table S4 from (19)). Precomputed gene expression changes following perturbation of the Wnt pathway by knocking down the repressor TCF3 were downloaded from a previous study (Table S2 from (20)). Gene symbols of both tables were modernized by converting outdated names to updated ones based on the a table from The Mouse Genome Nomenclature group (https://www.informatics.jax.org/mgihome/nomen/gene_name_initiative.shtml#completed).

#### Correlation and Variation among transcriptional protein signals

ChIP-seq peaks of 4 co-factors, MED1, BRD4, EP300, and CBP, were collapsed using bedtools merge. Peak-level signal was defined as ChIP-seq reads overlapping each collapsed peak calculated using bedtools intersect. Community-level signal was calculated by BOUQUET as above. Pairwise correlations of signal between proteins at both the peak and community level were measured by Spearman's correlation calculated by R function cor() with method='spearman' as parameter.

To investigate the variation among 3D-SEs across different co-factors, we compiled all communities that were called as 3D-SEs by their loading of at least one co-factor. This group of communities was hierarchically clustered based on their rankings across 4 co-factors using the heatmap.2() function from R package "gplots." In addition, Coefficients of Variation of per-community rankings across 4 co-factors were calculated using rowcvs() function from R package "Rfast2" (Supp. Data S21).

#### Gene ontology analysis

Gene ontology analysis was performed using the PanglaoDB_Augmented_2021 database within Enrichr (21–23) (https://maayanlab.cloud/Enrichr/) (Supp. Data S22). Genes associated 3D-SEs were independently used as input for unbiased enrichment analysis. For the comparison of genes associated with 3D-SEs in each cell type, the p-values for the top 10 ranked terms for each respective cell type were shown: mESCs, cardiomyocytes, SST-neurons, thymocytes, and regulatory T cells. For the comparison of genes associated with 3D-SEs for each co-activator and H3K27ac, the p-values for the top 10 ranked terms for each mark were shown.

**Gene Set Enrichment Analysis (GSEA) analysis**

For mESC cell identity gene analysis across multiple co-factors, genes were pre-ranked based on their community co-factor signal. We performed Preranked GSEA (24, 25) using the desktop version of the tool (v4.3.3) available at https://www.gsea-msigdb.org/gsea/downloads.jsp (Supp. Data S23). mESC cell identity genes are defined as above using the "Embryonic Stem Cell" gene set from the PanglaoDB_Augmented_2021 database available on Enrichr (https://maayanlab.cloud/Enrichr/) (21–23, 26).

For the analysis of mESC cell identity genes across multiple loop-calling FDR thresholds (0.005, 0.01, 0.05), CRE communities were identified separately at each threshold, and genes were then pre-ranked according to the MED1 signal of their respective communities.

For cell identity gene analysis across mouse cell types, genes were pre-ranked based on their community H3K27ac HiChIP signal. Cell identity genes of each mouse cell type are defined as genes associated with cell type specific PanglaoDB Augmented 2021 terms as follows: "Embryonic stem cells" for ESCs; "Cardiomyocytes" for Cardiomyocytes; "Neurons", "GABAergic Neurons" and "Interneurons" for SST-neurons; "T cells" and "Thymocytes" for Thymocytes; "T regulatory cells" and "T cells" for Tregs.

For the signaling pathway analysis, the fold changes in gene expression between two experimental conditions for each perturbed signaling pathway were used as a ranked list. Here, only genes that were determined to be active in wild-type mESCs were kept (see above in the "Inferring active genes/promoters" methods section for further details). Three gene sets were tested against the ranked expression results of each perturbation: 1) genes in 3D-SEs with a signaling-TF-bound CRE; 2) genes in communities with a signaling-TF-bound CRE; 3) expressed genes whose TSS was genomically most proximal to a signaling TF peak, determined using bedtools with "closest -t all".

**Cell culture for single-cell sequencing libraries**

The mouse ESC line V6.5 (a gift from the Young lab) was maintained in 2i/LIF medium at 37°C with 5% $CO_2$ in a humidified incubator. The culture medium was made with DMEM-F12 (Gibco, Cat# 11320033) was supplemented with 0.5X B27 (Gibco, Cat# 17504001), 0.5X N2 (Gibco, Cat# 17502001), 0.5mM L-glutamine (Gibco, Cat# 25030081), 0.1 mM beta-mercaptoethanol (Sigma, Cat# M6250-10ML), 1X Pen-Strep (Gibco, Cat# 15140122), 0.5X nonessential amino acids (Gibco, Cat# 11140050), 1000U/ml ESGRO recombinant LIF (Millipore, Cat# ESG1107), 1 µM PD0325901 (REPROCELL, Cat# 04-0006-02), 3 µM CHIR99021 (REPROCELL, Cat# 04-0004-02). Cells were cultured on vessel treated with 0.2% gelatin (Sigma, Cat# G1393-100ML). For passaging, cells dissociated using StemProAccutase (Gibco, Cat# A11105-01) and plated at a density of 30,000/cm$^2$.

#### Single-cell RNA-seq

##### scRNA-seq library preparation

Cell cultures at the log phase of growth were collected and checked for cell number and viability using TC20 automated counter (Bio-Rad, Cat# 1450102). Samples with >90 % viability were processed in order to prepare single cell suspensions and diluted into the target numbers plus the multiplet rate (1000 cells plus 0.8 % per library). Single cell RNA sequencing libraries were generated using the Chromium Next GEM Single Cell 5′ Kit v2 (10X Genomics, Cat# PN-1000263), following the manufacturer's manual. Briefly, single cell suspensions were loaded on the chromium controller to capture single cell gel beads and then used to perform in-droplet barcoding reactions followed by the amplification of full-length cDNAs. Sequencing libraries were constructed by sequential ligation reactions of kit-provided adaptor oligos and then dual indexed before enzymatically fragmenting the cDNAs. The resulting cDNA libraries were purified with solid-phase reversible immobilization SPRI beads (Beckman, Cat# B23318) and checked on a BioAnalyzer High sensitivity chip (Agilent, Cat# 5067-4626) and a Qubit fluorometer (InVitrogen, Cat# Q33238) for DNA concentrations and size distributions, respectively.

##### scRNA-seq sequencing

The libraries were sequenced on an Illumina NextSeq500 using the parameters recommended by 10X Genomics. For the run, the parameters "read1 26 base pair (bp) x i7 10 bp x i5 10 bp x read2 90 bp, 150-cycle High Output kit" were used to achieve reads depths of >50K per cell.

##### scRNA-seq data analysis

Cell Ranger v6.0.0 (10X Genomics) was used to process raw scRNA-seq sequencing data. The raw sequencing data from the Illumina sequencer were de-multiplexed. After converting Illumina raw basecall files (BCL) to FASTQ format using the command 'cellranger mkfastq,' sequencing reads were aligned to the mm10 transcriptome using the STAR aligner (27). The expression of each gene in each cell was quantified as the number of Unique Molecular Identifiers (UMI) for each gene per cell-specific cellular barcode. With ~61.8 million reads sequenced from mESCs, we obtained data from 849 cells that passed quality control steps implemented in Cell Ranger. In total, 67.2% of the total reads confidently mapped to the transcriptome. Reads that were confidently mapped to the transcriptome were used to generate gene-barcode matrices based on Chromium cellular barcodes. Filtered gene-barcode matrices containing cellular barcodes only were used for downstream analysis.

We carried out analyses of processed scRNA-seq data in R v4.0.2 with Seurat v4.1.0. As a further step of quality control, we filtered out genes expressed in <3 cells and cells with <200 or >8,000 detected genes. Additionally, cells with the percentage of mitochondrial genes >5% were filtered out. At the end, 847 cells survived this stringent quality control process. To normalize the raw read counts from the gene-barcode matrices, we employed a global-scaling normalization method "LogNormalize" that normalizes the gene expression measurements for each cell by the total expression, multiplies this by a scale factor of 10000, and log-transforms the result.

To examine transcriptome heterogeneity and attempt to find distinct cell clusters, we performed principal component analysis (PCA) to reduce data dimensionality using RunPCA command of Seurat. We first identified genes with highly variable expression across single cells using "VST" method. The normalized single-cell expression values of the top 2000 most variable genes were used. We selected the top 30 significant principal components using a permutation-based test and heuristic methods implemented in Seurat. The selected PCA loadings were used as input for graph-based cell clustering and as input for Uniform Manifold Approximation and Projection (UMAP) for reduction to two dimensions for visualization purposes.

**scRNA-seq based gene-expression correlation**

To measure pairwise gene expression association, we calculated Spearman correlation values of gene expression across the 847 high-quality cells from scRNA-seq data. Before correlation calculation, raw read counts of 15,396 genes, filtered as above were normalized using the "LogNormalize" method implemented in Seurat.

Due to the abundance of expression dropouts and overdispersion, the optimal measure of gene expression correlation based on scRNA-seq data is unclear (28). Therefore, we calculated another eight measures of gene expression association using the Dismay R package (29): Pearson correlation, Kendall rank correlation, Weighted rank correlation, Biweight midcorrelation, Canberra distance, Jaccard index, Cosine similarity, and Dice coefficient (Supp. Fig. 1G). P-value calculations are capped at the significance level of $p=2.23x10^{-308}$, which is the smallest positive double-precision floating point numbers in R v4.0.2

(.Machine$double.xmin). All such p-values that are impacted by this computational limit are reported as $p<2x10^{-300}$.

#### Single-cell ATAC-seq

##### scATAC-seq library preparation

Single-cell ATAC sequencing libraries were generated using the Chromium Next GEM Single Cell ATAC kit v2 (10X Genomics, Cat# PN-1000390), following the manufacturer's manual. Briefly, nuclei suspensions were prepared from single cells diluted in nuclei buffer. The transposition reaction of the nuclei was performed in the mixture of ATAC buffer and enzyme using a 30-minute incubation time. The transposed nuclei and barcoding reagents were loaded on the chromium controller to capture single-nucleus gel beads and then to perform in-droplet barcoding PCR. Sequencing libraries were constructed by the index PCR of the barcoded free DNA fragments. The resulting ATAC libraries were purified with solid-phase reversible immobilization SPRI beads (Beckman) and checked on a BioAnalyzer high-sensitivity chip (Agilent) and a Qubit fluorometer (InVitrogen) for DNA concentrations and size distributions, respectively.

##### scATAC-seq sequencing

The libraries were sequenced on an Illumina NextSeq500 using the parameters recommended by 10X Genomics. For the run, the parameters "read1 50 base pair (bp) x i7 8 bp x i5 16 bp x read2 60 bp, 150-cycle High Output kit" were used to achieve reads depths of >50K per nuclei.

##### scATAC-seq data analysis

We used the Cell Ranger ATAC pipeline v2.1.0 (10X Genomics) with the default parameters to analyze raw reads from mESC scATAC-seq including read de-multiplexing, read filtering and alignment, barcode counting, identification of transposase cut sites, cell calling, and count matrix generation. The raw sequencing data from the Illumina sequencer were de-multiplexed and converted to FASTQ format using command "cellranger-atac mkfastq." Reads were aligned to the mm10 mouse reference genome (version 2020-A-2.0.0) provided by 10X Genomics using the command "cellranger-atac count". The resulting peak-by-cell count matrices, fragment file and single cell metadata were further processed using R package Signac v1.11.0 (30) under R v4.3.1. To filter out low-quality cells, we computed the following quality control metrics using Signac: 1) number of reads in peaks, 2) percentage of reads in peaks, 3) transcription start site (TSS) enrichment score (TSSEnrichment()), and 4) nucleosome signal (NucleosomeSignal()). Cells were filtered based on the following metrics: number of reads in peaks within the range of 3000-60000; percentage of reads in peaks > 40%; nucleosome signal < 3; TSS enrichment score > 2. 3,171 cells survived this stringent quality control process. For cell clustering and basic analysis, we used peaks defined using Signac. We normalized peak counts by term frequency–inverse document frequency (TF-IDF) method using the Signac functions RunTFIDF(). We kept all features for downstream dimensional reduction by using FindTopFeatures with min.cutoff = ‘q0’. To reduce the dimensionality of data, we performed singular value decomposition (SVD) on the TD-IDF matrix using function RunSVD(). The resulting top latent semantic indexing (LSI) component was discarded as it correlated strongly with sequencing depth, and components 2-30 were used for downstream analysis, including further dimensionality reduction by uniform manifold approximation (UMAP) for data visualization (RunUMAP()) and neighborhood graph-based clustering (FindNeighbors() and FindClusters()). A pseudobulked scATAC-seq coverage file was generated by quantifying the genome-wide coverage of scATAC-seq reads from all cells using “bedtools genomecov” with default parameters and converting the output file to bigwig format using the tool “bedGraphToBigWig” (31).

#### Enhancer deletion analysis

##### Cell culture

A parental mouse Embryonic Stem Cell line (mESC) V6.5 and super-enhancer-deletion line V6.5_dMir290e were gifts provided by Dr. Young at the Whitehead Institute and were described in (32). V6.5 cells are male cells derived from a C57BL/6(F) x 129/sv(M) cross. All cell lines were cultured in 2i + LIF medium as described in the previous articles (11). Briefly, 967.5 mL DMEM/F12 (GIBCO, Cat# 11320033) was supplemented with 5 mL N2 supplement (GIBCO, Cat# 17502001), 10 mL B27 supplement (GIBCO, Cat# 17504001), 0.5 mM L-glutamine (GIBCO, Cat# 25030081), 0.5X non-essential amino acids (GIBCO, Cat# 11140050), 100 U/mL Penicillin-Streptomycin (GIBCO, Cat# 15140122), 0.1 mM b-mercaptoethanol (Sigma, Cat# M6250-10ML), 1 uM PD0325901 (REPROCELL, Cat# 04-0006-02), 3 µM CHIR99021 (REPROCELL, Cat# 04-0004-02), and 1000 U/mL ESGRO recombinant LIF (Millipore, Cat# ESG1107). Cells were negative for mycoplasma tested through the beginning to the end of culture, using the MycoAlert Mycoplasma Detection Kit (Lonza, Cat# LT07-318).

##### RNA-seq library preparation

RNA samples were isolated using AllPrep DNA/RNA Mini Kit (QIAGEN, Cat# 80204). RNA-seq libraries were prepared using KAPA RNA HyperPrep Kit with RiboErase (HRM) (Roche, Cat# KK8651). Briefly, 1 µg of RNA of each sample was mixed with 2 µl of ERCC ExFold spike-in RNA mixes (Invitrogen, Cat# 4456739) based on the RNA ratio recommended by the protocol, rRNA were depleted by hybridization of complementary DNA oligonucleotides and treatment with RNase and DNase. RNA samples were fragmented, and double stranded cDNA was generated by two steps of cDNA synthesis. Illumina TruSeq DNA UD Indexes (IDT, Cat# 20040870) were used to make libraries. Libraries were amplified using KAPA HiFi HotStart ReadyMix (Roche, Cat# KK2601) and Library Amplification Primer

Mix (Roche, Cat# 07958994001), followed by Qubit dsDNA quantification (Invitrogen, Cat# Q32853), and 2100 BioAnalyzer D1000 kit (Agilent, Cat#50671504) and Illumina MiSeq Nano v2 (Illumina, Cat# 15036714) QC.

#### Whole-genome sequencing

Genomic DNA was isolated from three mouse ES cell lines, V6.5, V6.5-meGFP-Brd4-meGFP and V6.5-meGFP-Med1, using the QIAGEN AllPrep DNA/RNA Mini kit (QIAGEN, Cat# 80204) and quantified using a Qubit fluorometer with the dsDNA BR kit (Thermo Fisher Scientific, Cat# Q32853). High-molecular-weight DNA was used for library preparation. Whole-genome sequencing libraries were prepared using the KAPA HyperPrep Library Preparation kit (Roche, Ref# KR0961) according to the manufacturer's instructions. Briefly, 500 ng of genomic DNA of each cell line was sheared to an average insert size of approximately 350 bp using a Covaris LE220, followed by end repair, A-tailing, and ligation of adapters from ITD for Illumina TrueSeq DNA UDI v2 (Illumina, Ref# 20042113). Adapter-ligated DNA was purified using AmPure XP beads (Beckman Coulter, Ref# A63881) and amplified by 3 cycles of PCR. Amplified libraries were assessed for size distribution and concentration using HS D1000 Screen Tape (Agilent, part# 5067) on a TapeStation system (Agilent, part# 4200) prior to sequencing. Further QC of libraries was performed using Illumina MiSeq Nano kits. Libraries were pooled at equimolar concentrations and loaded according to Illumina's recommended loading concentrations. The final sequencing was performed on an Illumina NextSeq 2000 platform using a P1 flow cell with a 300-cycle kit, generating paired-end 150 bp, and 25M PF reads for each sample. Base calling and demultiplexing were carried out using Illumina's onboard software.

#### Whole-genome sequencing analysis

Raw FASTQ reads were aligned to the mm10 revision of the mouse reference genome using bowtie v1.2.2 in single-end mode with parameters -k 1 -m 1 -n 2 and -l set to the read length. Coverage in 50 bp bins was computed with MACS v1.4.1 with parameters -w -S –shiftsize=200 –space=50 –nomodel and converted to bigWig format using wigToBigWig v4. Genome-wide coverage was visualized using IGV v2.19.7 (33).

#### Microscopy

##### Probe design, probe labeling, and antibodies used

For DNA fluorescence in situ hybridization (FISH), purified mouse BAC DNAs for *Sptlc2* (chr12:87239078-87411743 and chr12:87395112-87476186, mm10) were labeled with a red-dUTP (AF594, Molecular Probes) by nick translation. Purified mouse BAC DNA for *Esrrb* (chr12:86340459-86543664, mm10) was labeled with a green-dUTP (SeeBright 496, Enzo) by nick translation. TMED10 RP23-234G7 BAC DNA (chr12:85,337,154-85,480,126, mm10) was labeled with a green-dUTP

(SB496, Enzo). Chr12 RP24- 352N21 BAC DNA (chr12:30,177,561-30,315,451, mm10) was labeled with a green-dUTP (SB496, Enzo) by nick translation. BRD4 was tagged using AF647 Anti-Brd4 antibody (Abcam). For RNA FISH, probes were designed against introns of *Sptlc2* and *Esrrb* as detailed in Supplementary Data S26. *Sptcl2* nascent RNA probes were labeled with Cy3 fluorescent dye, and *Esrrb* nascent RNA probes were labeled with Cy5 fluorescent dye.

**DNA FISH cell culture, slide preparation, DNA probe hybridization, and antibody application**

250,000 single-cell mouse ESC cells V6.5 (4-5-23, a gift provided by Dr. Young at the Whitehead Institute) were applied to glass slides by cytocentrifugation. Slides were fixed in 1% PFA in PBS for 5 minutes followed by 1% PFA in PBS plus 0.05% Igepal for 5 minutes, and stored in 70% ETOH at -20°C. The slides were rinsed in PBS and blocked in 200 µl blocking buffer (2X SSC and 1% BSA) and then incubated with an AF647 Anti-Brd4 antibody (Abcam) diluted at 1:100 in antibody diluent (2X SSC and 1% BSA) for 45 minutes at RT in the dark. Slides were then washed in PBS, placed in 4% PFA for 2 minutes at RT, rinsed in PBS, and dehydrated in 70%, 80%, 100% ETOH for 2 minutes each. The slides were then washed in 50% formamide and 2X SSC and mounted in Vectashield containing DAPI.-Cross linking with 4% PFA for 8 minutes was then performed to stabilize antibodies on proteins, followed by treatment in 0.2N HCL for 10 minutes on ice. Slides were then denatured in 70% formamide 2X SSC at 80°C for 10 minutes followed by ETOH dehydration series. Denatured DNA probes were then added in a solution containing mouse sheared cot DNA, 50% formamide, 2X SSC, and 10% dextran and hybridized at 37 °C overnight. Slides were then washed in 50% formamide, 2X SSC at 37 °C and mounted in Vectashield containing DAPI and imaged.

**RNA FISH cell culture, slide preparation, and RNA probe hybridization**

RNA FISh experiments were carried out in two separate V6.5 mESC cell lines: V6.5 mESCs containing MED1 endogenously tagged with GFP or BRD4 endogenously tagged with GFP (mESC-V6.5-MED1-GFP, mESC-V6.5-BRD4-GFP cells were a gift provided by Dr. Young at the Whitehead Institute). Single cell suspensions made from each cell line and were used to make cytospin slides. For each cytospin slide single-cell cells were applied to glass slides by cytocentrifugation. Slides were fixed in 1% PFA in PBS for 5 minutes followed by 1% PFA in PBS plus 0.05% Igepal for 5 minutes, and stored in 70% ETOH at -20°C. For the assay, a slide was briefly rinsed in PBS, dehydrated in a 3-part ETOH dehydration series consisting of 70%, 80%, and 100% ETOH each for 2 minutes, and then stained with Vectashield mounting medium containing DAPI and imaged for endogenous GFP. After imaging, the coverslip was removed and the slide was placed in 4% PFA for 2 minutes, then in 70%, 80%, and 100% ETOH each for 2 minutes. The slide was hybridized to 20 µl of probe (vial 1) in hybridization buffer containing 50% formamide, 10% dextran sulfate, 2X SSC, and 0.02%Tween and hybridized overnight at 42°C followed by washing in gradient washes (A, B, B, C). 20µl of denatured probe (vial 2) in same hybridization buffer as above was applied to the slide and hybridized for 3 hours at 40°C followed by washing in gradient washes (A, B, B, C). 20µl of the detection probe (vial 3) in the same hybridization buffer was applied to the slide and hybridized for 3 hours at 40°C followed by washing in gradient washes (A, B, B, C). After this last wash, the slide was dehydrated in a 3-part ETOH dehydration series consisting of 70%, 80%, 100% ETOH each for 2 minutes, and then stained with Vectashield mounting medium containing DAPI and imaged.

**Image analysis and scoring**

Imaging was performed in 3D using a widefield fluorescence microscope (Nikon Eclipse Ti2) equipped with Nikon Nis Elements version 6.02 software and Autoquant 3D deconvolution. Images were deconvolved using the Richardson-Lucy with number of rounds set to auto-select. Images were acquired using 0.1-micron (0.1 µm) plane spacing and a 60X planapochromatic objective with a numerical aperture of 1.4.

The captured images were analyzed by in-house Z-Fisher software, which is maintained by the Cytogenetics Shared Resource at St. Jude Children's Research Hospital in Memphis, TN. DNA FISH with immunofluorescence and RNA FISH and endogenous fluorescence images were auto aligned, and nuclei were gated by the software. Maximum intensity projections (MIP) were automatically generated for feature identification. Manual nuclei gating was performed on the DAPI MIP to remove and re-gate any erroneous software generated nuclei gates. Regions of Interest (ROI) were independently selected for 3 channels (False colored: Esrrb-Magenta, Sptlc2-Yellow, BRD4 IF-Green, MED1-GFP-Green, BRD4-GFP-Green). Colocalization distance thresholds were set to 0.40 µm. Distance calculations in the X, Y, and Z dimensions were autogenerated and any ROI that were within the threshold distance were tallied for reporting.

#### Data visualization

We used the UCSC Browser (31) (*http://genome.ucsc.edu)* and ProteinPaint ([https://proteinpaint.stjude.org/](https://proteinpaint.stjude.org/)) (34) to display various types of genome sequence data using the following Track Formats: hic for HiChIP interactions; bigInteract for HiChIP loops; bigGenePhred for gene annotation; bigwig (35) for HiChIP, ChIP-seq, pseudobulked scATAC and RNA-Seq coverage; bigBed (35) for genomic locations of enhancers, promoters, INs, and TADs. We used basic R for box-and-whisker plots, the R package "ggplot2" for violin plots, scatterplots and binned box-and-whisker plots; the R package "Vennerable" for Chow-Ruskey diagrams; the R package "igraph" for network diagrams; the R package "Seurat" for scRNA-Seq Uniform Manifold Approximation and Projection (UMAP) plots; the R package "Signac" for scATAC-Seq UMAP plots; and Adobe Illustrator and

BioRender.com for heatmap and schematic generation. R package "Reshape2" and "ggplot2" for spearman correlation heatmaps; R package "gplots" for hierarchical heatmap of community rankings. P-value calculations are capped at the significance level of $p=2.23x10^{-308}$, which is the smallest positive double-precision floating point numbers in R v4.0.2 (.Machine$double.xmin). All such p-values that are impacted by this computational limit are reported as $p<2x10^{-300}$.

49 Spadina Ave. Suite 200
Toronto ON M5V 2J1 Canada
www.biorender.com

### Confirmation of Publication and Licensing Rights - Open Access

**January 14th, 2026**

| | |
|---|---|
| ***Subscription Type:*** | *Institution - Academic* |
| ***Agreement number:*** | *OV298MOCPO* |
| ***Publisher Name:*** | *Nucleic Acids Research* |
| ***Figure Title:*** | *Figure 4: 3D-SEs colocalize with putative co-activator condensates.* |
| ***Citation to Use:*** | *Created in BioRender. Maher, K. (2026) https://BioRender.com/fghhosi* |

To whom this may concern,

This document ("Confirmation") hereby confirms that Science Suite Inc. dba BioRender ("BioRender") has granted the following BioRender user: Kelsey Maher ("User") a BioRender Academic Publication License in accordance with BioRender's Terms of Service and Academic License Terms ("License Terms") to permit such User to do the following on the condition that all requirements in this Confirmation are met:

1) publish their Completed Graphics created in the BioRender Services containing both User Content and BioRender Content (as both are defined in the License Terms) in publications (journals, textbooks, websites, etc.); and
2) sublicense such Completed Graphics under "open access" publication sublicensing models such as CC-BY 4.0 and more restrictive models, so long as the conditions set forth herein are fully met.

Requirements of User:

1) All Completed Graphics to be published in any publication (journals, textbooks, websites, etc.) must be accompanied by the following citation either as a caption, footnote or reference for each figure that includes a Completed Graphic:
"Created in BioRender. Maher, K. (2026) https://BioRender.com/fghhosi ".
2) All terms of the License Terms including all Prohibited Uses are fully complied with. E.g. For Academic License Users, no commercial uses (beyond publication in journals, textbooks or websites) are permitted without obtaining or switching to a BioRender Industry Plan.
3) A Reader (defined below) may request that the User allow their figure to be a public template for Readers to view, copy, and modify the figure. It is up to the User to determine what level of access to grant.

Open-Access Journal Readers:

Open-Access journal readers ("Reader") who wish to view and/or re-use a particular Completed Graphic in an Open-Access journal subject to CC-BY sublicensing may do so by clicking on the URL link in the applicable citation for the subject Completed Graphic.

The re-use/modification options below are available after the Reader requests the User to adapt their figure as a BioRender template and the User has granted such access.

1) View-Only/Free Plan Use: A Reader who wishes to only view the Completed Graphic may do so in the BioRender Services as either a BioRender Free Plan user or simply as a viewer. By becoming a BioRender Free Plan user, the Reader may view, modify and re-use the Completed Graphic as permitted under BioRender's Basic License Terms (e.g. personal use only, no publishing or commercial use permitted).
2) Re-Use/Publish with No Modifications: For any re-use and re-publication of a Completed Graphic with no modification(s) to the Completed Graphic made by the Reader, a Reader may do so by citing the original author using the citation noted above with the Completed Graphic. The Reader must also comply with the underlying License Terms which apply to the Completed Graphic as noted above (e.g. no commercial use for Academic License).
3) Re-Use/Publish with Modifications: For any re-use and re-publication of a Completed Graphic with a modification(s) made by the Reader, the Reader may do so by becoming a BioRender user themselves under either an Academic or Industry Plan, citing the original author using the citation noted above with the Completed Graphic and complying with the applicable License Terms.

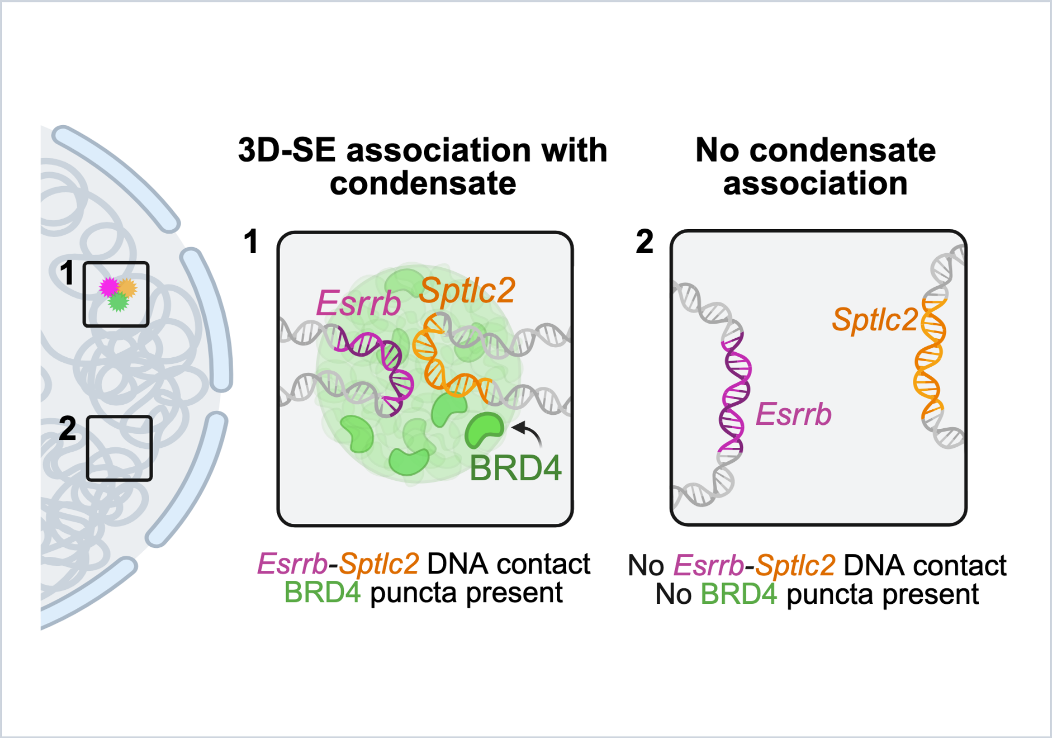


*For any questions regarding this document, or other questions about publishing with BioRender, please refer to our BioRender Publication Guide, or contact BioRender Support at.*

49 Spadina Ave. Suite 200
Toronto ON M5V 2J1 Canada
www.biorender.com

### Confirmation of Publication and Licensing Rights - Open Access

**July 1st, 2025**

| | |
|---|---|
| ***Subscription Type:*** | *Institution - Academic* |
| ***Agreement number:*** | *RO28GI61K3* |
| ***Publisher Name:*** | *Nucleic Acids Research* |
| ***Figure Title:*** | *Figure 4: 3D-SEs colocalize with putative co-activator condensates.* |
| ***Citation to Use:*** | *Created in BioRender. Maher, K. (2025) https://BioRender.com/ycwql9z* |

To whom this may concern,

This document ("Confirmation") hereby confirms that Science Suite Inc. dba BioRender ("BioRender") has granted the following BioRender user: Kelsey Maher ("User") a BioRender Academic Publication License in accordance with BioRender's Terms of Service and Academic License Terms ("License Terms") to permit such User to do the following on the condition that all requirements in this Confirmation are met:

1) publish their Completed Graphics created in the BioRender Services containing both User Content and BioRender Content (as both are defined in the License Terms) in publications (journals, textbooks, websites, etc.); and
2) sublicense such Completed Graphics under "open access" publication sublicensing models such as CC-BY 4.0 and more restrictive models, so long as the conditions set forth herein are fully met.

Requirements of User:

1) All Completed Graphics to be published in any publication (journals, textbooks, websites, etc.) must be accompanied by the following citation either as a caption, footnote or reference for each figure that includes a Completed Graphic:
   "Created in BioRender. Maher, K. (2025) https://BioRender.com/ycwql9z ".
2) All terms of the License Terms including all Prohibited Uses are fully complied with. E.g. For Academic License Users, no commercial uses (beyond publication in journals, textbooks or websites) are permitted without obtaining or switching to a BioRender Industry Plan.
3) A Reader (defined below) may request that the User allow their figure to be a public template for Readers to view, copy, and modify the figure. It is up to the User to determine what level of access to grant.

Open-Access Journal Readers:

Open-Access journal readers ("Reader") who wish to view and/or re-use a particular Completed Graphic in an Open-Access journal subject to CC-BY sublicensing may do so by clicking on the URL link in the applicable citation for the subject Completed Graphic.

The re-use/modification options below are available after the Reader requests the User to adapt their figure as a BioRender template and the User has granted such access.

1) View-Only/Free Plan Use: A Reader who wishes to only view the Completed Graphic may do so in the BioRender Services as either a BioRender Free Plan user or simply as a viewer. By becoming a BioRender Free Plan user, the Reader may view, modify and re-use the Completed Graphic as permitted under BioRender's Basic License Terms (e.g. personal use only, no publishing or commercial use permitted).
2) Re-Use/Publish with No Modifications: For any re-use and re-publication of a Completed Graphic with no modification(s) to the Completed Graphic made by the Reader, a Reader may do so by citing the original author using the citation noted above with the Completed Graphic. The Reader must also comply with the underlying License Terms which apply to the Completed Graphic as noted above (e.g. no commercial use for Academic License).
3) Re-Use/Publish with Modifications: For any re-use and re-publication of a Completed Graphic with a modification(s) made by the Reader, the Reader may do so by becoming a BioRender user themselves under either an Academic or Industry Plan, citing the original author using the citation noted above with the Completed Graphic and complying with the applicable License Terms.

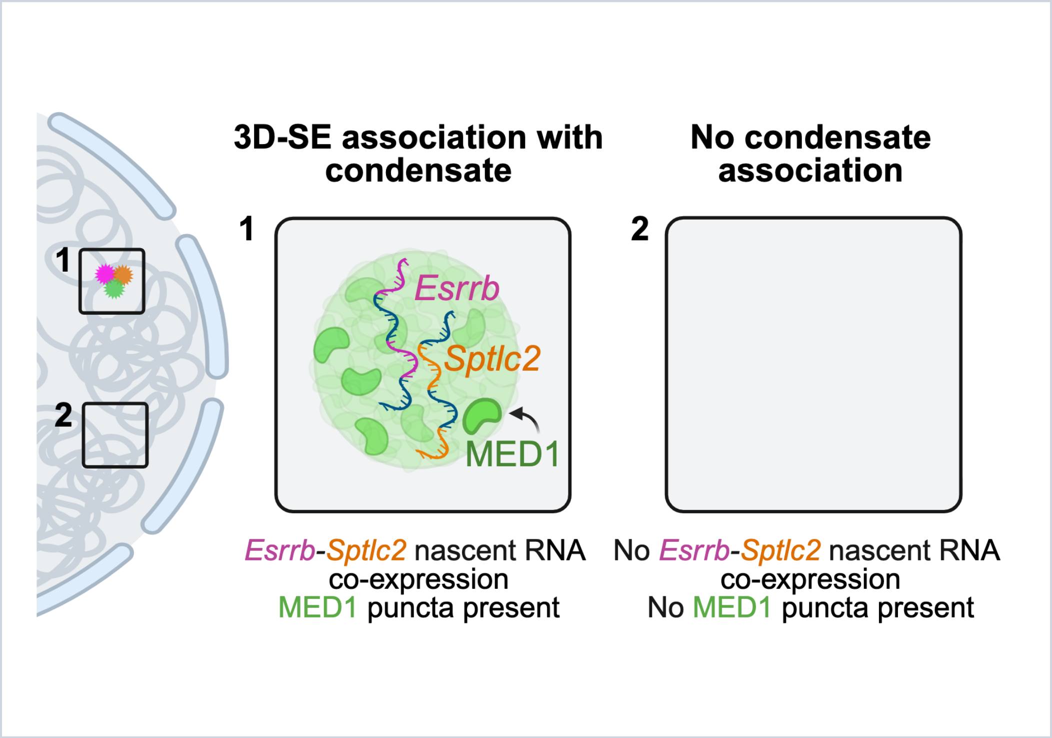


*For any questions regarding this document, or other questions about publishing with BioRender, please refer to our BioRender Publication Guide, or contact BioRender Support at.*

49 Spadina Ave. Suite 200
Toronto ON M5V 2J1 Canada
www.biorender.com

### Confirmation of Publication and Licensing Rights - Open Access

**July 1st, 2025**

| | |
|---|---|
| ***Subscription Type:*** | *Institution - Academic* |
| ***Agreement number:*** | *XA28GILJX5* |
| ***Publisher Name:*** | *Nucleic Acids Research* |
| ***Figure Title:*** | *Figure S9: Fluorescence microscopy of co-localized 3D-SE components.* |
| ***Citation to Use:*** | *Created in BioRender. Maher, K. (2025) https://BioRender.com/jkj4wow* |

To whom this may concern,

This document ("Confirmation") hereby confirms that Science Suite Inc. dba BioRender ("BioRender") has granted the following BioRender user: Kelsey Maher ("User") a BioRender Academic Publication License in accordance with BioRender's Terms of Service and Academic License Terms ("License Terms") to permit such User to do the following on the condition that all requirements in this Confirmation are met:

1) publish their Completed Graphics created in the BioRender Services containing both User Content and BioRender Content (as both are defined in the License Terms) in publications (journals, textbooks, websites, etc.); and
2) sublicense such Completed Graphics under "open access" publication sublicensing models such as CC-BY 4.0 and more restrictive models, so long as the conditions set forth herein are fully met.

Requirements of User:

1) All Completed Graphics to be published in any publication (journals, textbooks, websites, etc.) must be accompanied by the following citation either as a caption, footnote or reference for each figure that includes a Completed Graphic:
   "Created in BioRender. Maher, K. (2025) https://BioRender.com/jkj4wow ".
2) All terms of the License Terms including all Prohibited Uses are fully complied with. E.g. For Academic License Users, no commercial uses (beyond publication in journals, textbooks or websites) are permitted without obtaining or switching to a BioRender Industry Plan.
3) A Reader (defined below) may request that the User allow their figure to be a public template for Readers to view, copy, and modify the figure. It is up to the User to determine what level of access to grant.

Open-Access Journal Readers:

Open-Access journal readers ("Reader") who wish to view and/or re-use a particular Completed Graphic in an Open-Access journal subject to CC-BY sublicensing may do so by clicking on the URL link in the applicable citation for the subject Completed Graphic.

The re-use/modification options below are available after the Reader requests the User to adapt their figure as a BioRender template and the User has granted such access.

1) View-Only/Free Plan Use: A Reader who wishes to only view the Completed Graphic may do so in the BioRender Services as either a BioRender Free Plan user or simply as a viewer. By becoming a BioRender Free Plan user, the Reader may view, modify and re-use the Completed Graphic as permitted under BioRender's Basic License Terms (e.g. personal use only, no publishing or commercial use permitted).
2) Re-Use/Publish with No Modifications: For any re-use and re-publication of a Completed Graphic with no modification(s) to the Completed Graphic made by the Reader, a Reader may do so by citing the original author using the citation noted above with the Completed Graphic. The Reader must also comply with the underlying License Terms which apply to the Completed Graphic as noted above (e.g. no commercial use for Academic License).
3) Re-Use/Publish with Modifications: For any re-use and re-publication of a Completed Graphic with a modification(s) made by the Reader, the Reader may do so by becoming a BioRender user themselves under either an Academic or Industry Plan, citing the original author using the citation noted above with the Completed Graphic and complying with the applicable License Terms.

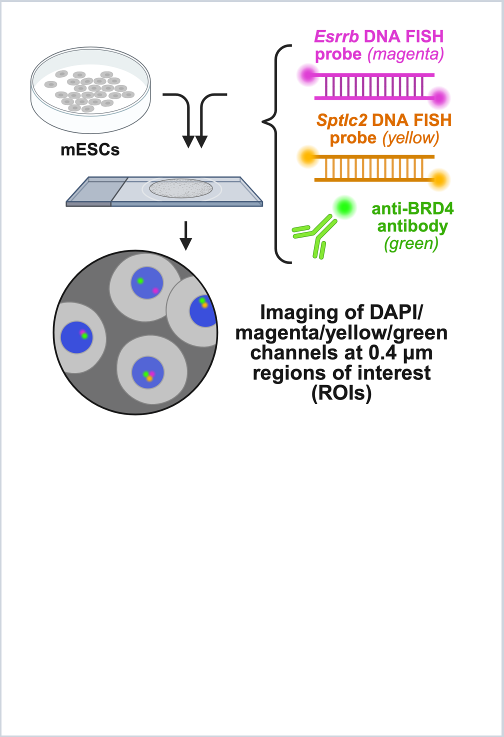


*For any questions regarding this document, or other questions about publishing with BioRender, please refer to our BioRender Publication Guide, or contact BioRender Support at.*

49 Spadina Ave. Suite 200
Toronto ON M5V 2J1 Canada
www.biorender.com

### Confirmation of Publication and Licensing Rights - Open Access

**July 1st, 2025**

***Subscription Type:*** *Institution - Academic*
***Agreement number:*** *HG28GIARRL*
***Publisher Name:*** *Nucleic Acids Research*

***Figure Title:*** *Figure S9: Fluorescence microscopy of co-localized 3D-SE components.*

***Citation to Use:*** *Created in BioRender. Maher, K. (2025) https://BioRender.com/b9h2fpm*

To whom this may concern,

This document ("Confirmation") hereby confirms that Science Suite Inc. dba BioRender ("BioRender") has granted the following BioRender user: Kelsey Maher ("User") a BioRender Academic Publication License in accordance with BioRender's Terms of Service and Academic License Terms ("License Terms") to permit such User to do the following on the condition that all requirements in this Confirmation are met:

1) publish their Completed Graphics created in the BioRender Services containing both User Content and BioRender Content (as both are defined in the License Terms) in publications (journals, textbooks, websites, etc.); and
2) sublicense such Completed Graphics under "open access" publication sublicensing models such as CC-BY 4.0 and more restrictive models, so long as the conditions set forth herein are fully met.

Requirements of User:

1) All Completed Graphics to be published in any publication (journals, textbooks, websites, etc.) must be accompanied by the following citation either as a caption, footnote or reference for each figure that includes a Completed Graphic:
"Created in BioRender. Maher, K. (2025) https://BioRender.com/b9h2fpm ".
2) All terms of the License Terms including all Prohibited Uses are fully complied with. E.g. For Academic License Users, no commercial uses (beyond publication in journals, textbooks or websites) are permitted without obtaining or switching to a BioRender Industry Plan.
3) A Reader (defined below) may request that the User allow their figure to be a public template for Readers to view, copy, and modify the figure. It is up to the User to determine what level of access to grant.

Open-Access Journal Readers:

Open-Access journal readers ("Reader") who wish to view and/or re-use a particular Completed Graphic in an Open-Access journal subject to CC-BY sublicensing may do so by clicking on the URL link in the applicable citation for the subject Completed Graphic.

The re-use/modification options below are available after the Reader requests the User to adapt their figure as a BioRender template and the User has granted such access.

1) View-Only/Free Plan Use: A Reader who wishes to only view the Completed Graphic may do so in the BioRender Services as either a BioRender Free Plan user or simply as a viewer. By becoming a BioRender Free Plan user, the Reader may view, modify and re-use the Completed Graphic as permitted under BioRender's Basic License Terms (e.g. personal use only, no publishing or commercial use permitted).
2) Re-Use/Publish with No Modifications: For any re-use and re-publication of a Completed Graphic with no modification(s) to the Completed Graphic made by the Reader, a Reader may do so by citing the original author using the citation noted above with the Completed Graphic. The Reader must also comply with the underlying License Terms which apply to the Completed Graphic as noted above (e.g. no commercial use for Academic License).
3) Re-Use/Publish with Modifications: For any re-use and re-publication of a Completed Graphic with a modification(s) made by the Reader, the Reader may do so by becoming a BioRender user themselves under either an Academic or Industry Plan, citing the original author using the citation noted above with the Completed Graphic and complying with the applicable License Terms.

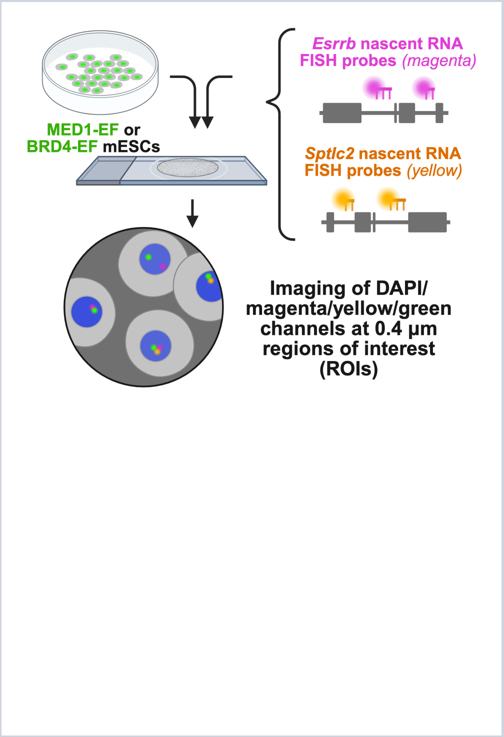


*For any questions regarding this document, or other questions about publishing with BioRender, please refer to our BioRender Publication Guide, or contact BioRender Support at.*
